# Distorted tonotopy severely degrades neural representations of natural speech in noise following acoustic trauma

**DOI:** 10.1101/2021.04.22.440950

**Authors:** Satyabrata Parida, Michael G. Heinz

**Affiliations:** Weldon School of Biomedical Engineering, Purdue University, West Lafayette, IN, 47907 USA; Department of Speech, Language, and Hearing Sciences, Purdue University, West Lafayette, IN, 47907 USA

**Keywords:** Speech coding, noise-induced hearing loss, tonotopic coding, auditory nerve, suprathreshold deficits, speech audibility, temporal precision

## Abstract

Listeners with sensorineural hearing loss (SNHL) struggle to understand speech, especially in noise, despite audibility compensation. These real-world suprathreshold deficits are hypothesized to arise from degraded frequency tuning and reduced temporal-coding precision; however, peripheral neurophysiological studies testing these hypotheses have been largely limited to in-quiet artificial vowels. Here, we measured single auditory-nerve-fiber responses to a natural speech sentence in noise from anesthetized chinchillas with normal hearing (NH) or noise-induced hearing loss (NIHL). Our results demonstrate that temporal precision was not degraded, and broader tuning was not the major factor affecting peripheral coding of natural speech in noise. Rather, the loss of cochlear tonotopy, a hallmark of normal hearing, had the most significant effects (both on vowels and consonants). Because distorted tonotopy varies in degree across etiologies (e.g., noise exposure, age), these results have important implications for understanding and treating individual differences in speech perception for people suffering from SNHL.

## INTRODUCTION

Individuals with SNHL demonstrate speech-perception deficits, especially in noise, which are often not resolved even with state-of-the-art hearing-aid strategies and noise-reduction algorithms (Lesica, 2018; McCormack and Fortnum, 2013). In fact, difficulty understanding speech in noise is the number-one complaint in audiology clinics (Chung, 2004), and can leave people with SNHL suffering from communication difficulties that impact their professional, social, and family lives, as well as their mental health (Dawes et al., 2015; Mener et al., 2013). Although audibility is a factor contributing to these difficulties (Phatak and Grant, 2014), it is clear that other suprathreshold deficits associated with dynamic spectrotemporal cues contribute as well, especially in noise (Festen and Plomp, 1983; Zeng and Turner, 1990). Unfortunately, neurophysiological studies of speech coding in the impaired auditory nerve (AN), the first neural site affected by cochlear components of SNHL (Trevino et al., 2019), have been primarily limited to synthetic vowels in quiet and have not included everyday sounds with natural dynamics.

The two most common suprathreshold factors hypothesized to underlie speech-perception deficits are degraded frequency selectivity and reduced temporal-coding precision. Broader tuning, often observed in listeners with SNHL (Glasberg and Moore, 1986), may limit the ability to resolve spectral components of speech and allow more background noise into auditory filters (Moore, 2007). Reduced perceptual frequency selectivity parallels the broader tuning observed in physiological responses following outer-hair-cell dysfunction (Liberman and Dodds, 1984a; Ruggero and Rich, 1991). Broadened frequency selectivity often correlates with degraded speech perception in noise (e.g., Festen and Plomp, 1983; Glasberg and Moore, 1989), with listeners with SNHL being more susceptible to noise than listeners with NH (Horst, 1987). A second suprathreshold deficit hypothesized to underlie degraded speech perception, especially in noise, is a loss of temporal-coding precision following SNHL (Halliday et al., 2019; Lorenzi et al., 2006; Moore, 2008). While one study has reported a decrease in AN phase locking following SNHL (Woolf et al., 1981), others have not found a degradation (Harrison and Evans, 1979; Kale and Heinz, 2010); however, these studies have been limited to laboratory stimuli (e.g., tones or modulated tones).

Another neural suprathreshold factor that may contribute to speech-coding degradations is the change in on-versus off-frequency sensitivity (i.e., reduced tip-to-tail ratio, TTR) observed in AN fibers following NIHL. This phenomenon, which results from a combination of reduced tip sensitivity (and associated loss of tuning) and hypersensitive tails, has been well characterized with tonal frequency-tuning curves (FTCs; Liberman and Dodds, 1984a). These distortions in tonotopic sensitivity have important implications for complex-sound processing but have only recently begun to be explored for nontonal stimuli such as broadband noise (Henry et al., 2016, 2019).

Several neurophysiological studies have investigated speech coding following SNHL; however, these studies have not probed natural speech coding in noise, for which listeners with SNHL struggle the most (Sayles and Heinz, 2017; Young, 2012). Studies of NH AN speech coding have predominantly used short-duration synthesized vowel- and consonant-like stimuli in quiet (e.g., Delgutte and Kiang, 1984b, 1984c; Sinex and Geisler, 1983; Young and Sachs, 1979), with the few exploring background noise limited to vowels (Delgutte and Kiang, 1984a; Sachs et al., 1983). A few studies of NH coding have used natural speech sentences but not in noise (Delgutte et al., 1998; Kiang and Moxon, 1974; Young, 2008). Speech-coding studies in hearing-impaired (HI) animals have been primarily limited to vowel-like stimuli in quiet (e.g., Miller et al., 1997; Schilling et al., 1998). Natural speech differs from synthetic speech tokens in its highly dynamic nature, with most information contained in its time-varying properties like formant transitions and spectrotemporal modulations (Elhilali, 2019; Elliott and Theunissen, 2009). The limited study of natural-speech coding is likely due to standard spike-train Fourier-based analyses (e.g., Young and Sachs, 1979) not being suitable for temporally varying stimuli (Parida et al., 2021).

Here, we used a natural speech sentence stimulus and collected AN-fiber spike trains from anesthetized chinchillas with either NH or NIHL. The sentence was also mixed with speech-shaped noise at perceptually relevant signal-to-noise ratios (SNRs). We analyzed dynamic formant transitions in vowel responses in noise using newly developed nonstationary analyses based on frequency demodulation of alternating-polarity peristimulus-time histograms (Parida et al., 2021), and analyzed onset and sustained responses of fricatives and stop consonants. Our results provide unique insight into the physiological suprathreshold mechanisms that do and do not contribute to degraded natural speech coding in noise, specifically highlighting the important role distorted tonotopy plays in increased noise susceptibility following SNHL. These findings have important implications for better understanding individual differences in speech perception in people with SNHL.

## RESULTS

### Chinchilla model of NIHL captures reduced audibility and degraded frequency selectivity

Mild-to-moderate hearing loss is the most prevalent degree among patients with hearing loss (Goman and Lin, 2016). To investigate the neural coding deficits these patients likely experience, we used an established noise-exposure protocol to induce mild-to-moderate SNHL (Kale and Heinz, 2010). Thresholds for auditory brainstem responses (ABRs) to tone bursts increased by ~20 dB following NIHL (Figure 1A, statistics in legends). Similarly, DPOAE levels decreased by ~15 dB (Figure 1B), indicating the presence of substantial outer-hair-cell damage. These electrophysiological changes indicate a mild-to-moderate permanent hearing-loss model (Clark, 1981).

**Figure 1.**
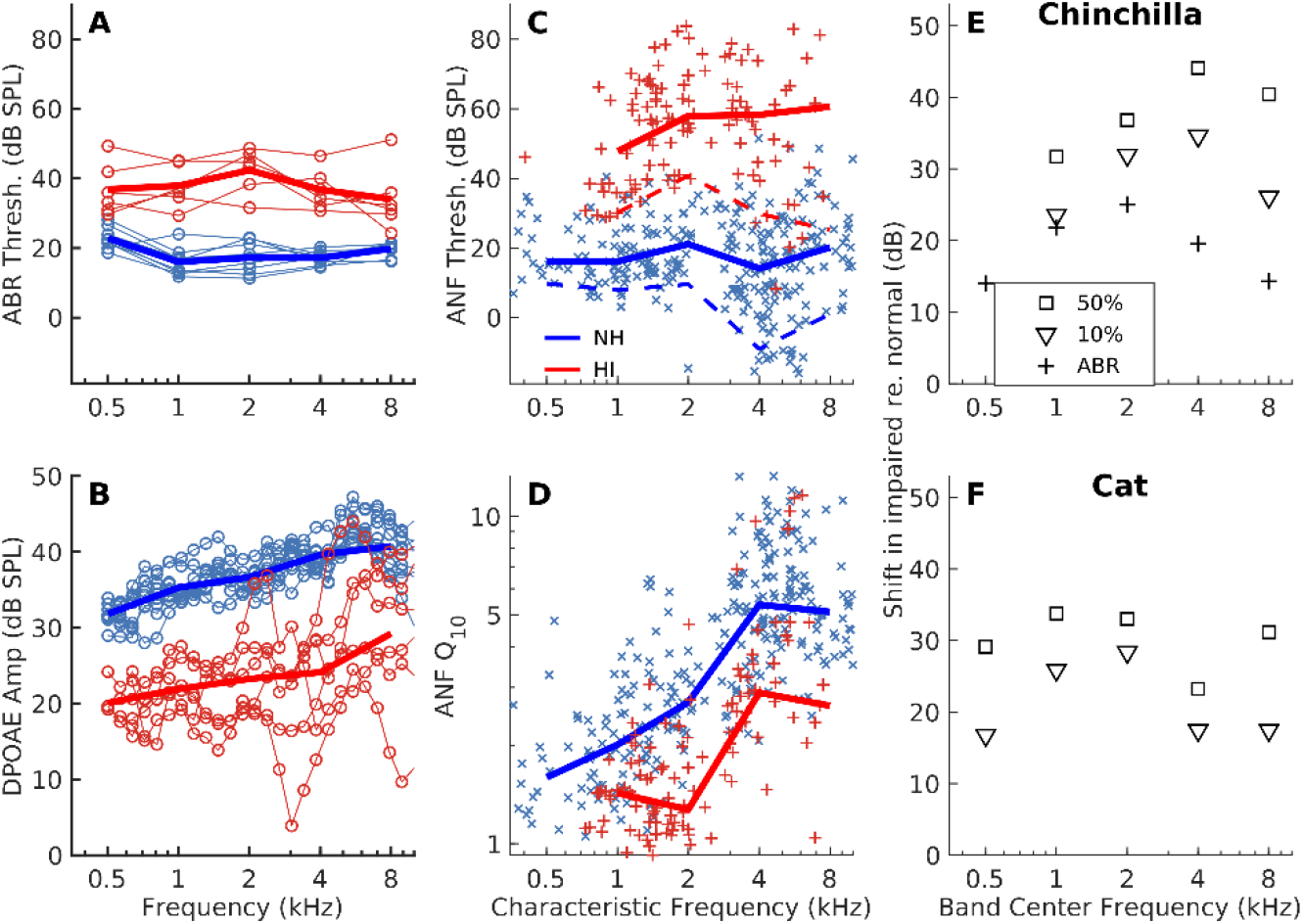
Chinchilla model of noise-induced hearing loss (NIHL) captures reduced audibility, broader frequency selectivity, and expanded AN-fiber threshold distribution. (A) Auditory-brainstem-response (ABR) thresholds for hearing-impaired (HI) chinchillas (red) were elevated by ~20 dB relative to normal hearing (NH, blue). Thin lines with symbols represent individual animals (n=9/6, NH/HI); thick lines represent group averages (main effect of group, F=276.4, p<2.2×10^−16^). (B) Distortion-product otoacoustic emission (DPOAE) amplitudes were reduced by ~15 dB (F=943.6, p<2×10^−16^). (C) AN-fiber thresholds were elevated for the HI group (F=646.36, p<2×10^−16^; n=286/119, NH/HI). Markers represent individual AN fibers; solid and dashed lines represent 50^th^ and 10^th^ percentiles, respectively, computed within octave bands for which there were ≥7 fibers in the group. (D) Q_10_ tuning-sharpness values were reduced for HI AN fibers (F=53.87, p=1.2×10^−12^). Thick lines represent octave-band averages. (E) Shifts in 50^th^ percentile chinchilla AN-fiber thresholds (squares) following NIHL were greater than shifts in the 10^th^ percentile (triangles). ABR-threshold shifts are shown with plusses. (F) Same format as E, but showing the same threshold-distribution expansion in a more extensive noise-exposed data set from cat (n=180/558, NH/HI), reanalyzed from Heinz et al. (2005).

Single-fiber thresholds were elevated by ~25-35 dB for the most-sensitive AN fibers in the population (Figures 1C and 1E). This threshold shift was accompanied by substantial loss of tuning as quantified by reductions in local Q10 values for individual-fiber FTCs (Figure 1D). These physiological effects were similar to previous results from our laboratory (Henry et al., 2016; Kale and Heinz, 2010).

### NIHL expands AN-fiber threshold distribution: Audiometric threshold shift underestimates average fiber-audibility deficit

Although audiometric thresholds likely relate to the most-sensitive AN fibers in the population, suprathreshold speech perception in complex environments likely requires integration across many fibers, not just the most sensitive ones (Bharadwaj et al., 2014). These effects can be assessed by characterizing changes to the AN-fiber threshold distribution. A traditional hypothesis, assuming the most sensitive fibers are the most vulnerable, predicts a compressed AN threshold distribution following NIHL (Moore et al., 1985; Ngan and May, 2001). Alternatively, an expanded distribution could lead to uncompensated audibility for most AN fibers if gain were based on the most sensitive fibers (e.g., using the audiogram). This effect could potentially contribute to poorer audibility of consonants compared to vowels despite similar audiometric compensation (Phatak and Grant, 2014; Zeng and Turner, 1990).

To evaluate these hypotheses, we estimated 10^th^ and 50^th^ percentiles for our NH and HI threshold distributions in octave-wide characteristic-frequency (CF) bands (Figure 1E). Our results showed a greater shift for the 50^th^ percentiles compared to the 10^th^ percentiles following NIHL for all bands. We also reanalyzed a more extensive previously published data set from cats (Heinz et al., 2005), which showed the same expansion (not compression) in AN-fiber threshold distribution following NIHL (Figure 1F). These consistent results suggest that any audiometric indicator (i.e., based on the most sensitive AN fibers) will underestimate the audibility deficits in many AN fibers across the population.

### Temporal-coding precision for natural speech was not degraded by NIHL

To test whether there was any degradation in the ability of AN fibers to precisely encode temporal information in response to natural speech, trial-to-trial precision was quantified using the Victor-Purpura (VP) distance metric (Victor and Purpura, 1996). Temporal precision is inversely related to VP distance, and was computed for a range of temporal resolutions using the time-shift cost parameter (*q*) of the VP analysis to span the syllabic, voice-pitch, and formant time scales of speech (Rosen, 1992). Temporal precision of natural-speech responses was not degraded following NIHL for any of the temporal resolutions considered (Figure 2). In fact, there was a small but significant increase in precision for all three time-scale conditions. This increase in precision may arise due to overrepresentation of lower-frequency information associated with distorted tonotopy, where synchrony is stronger than higher frequencies. Overall, these data provide no evidence for a degradation in the fundamental ability of AN fibers to precisely phaselock to the temporal features in natural speech following NIHL.

**Figure 2.**
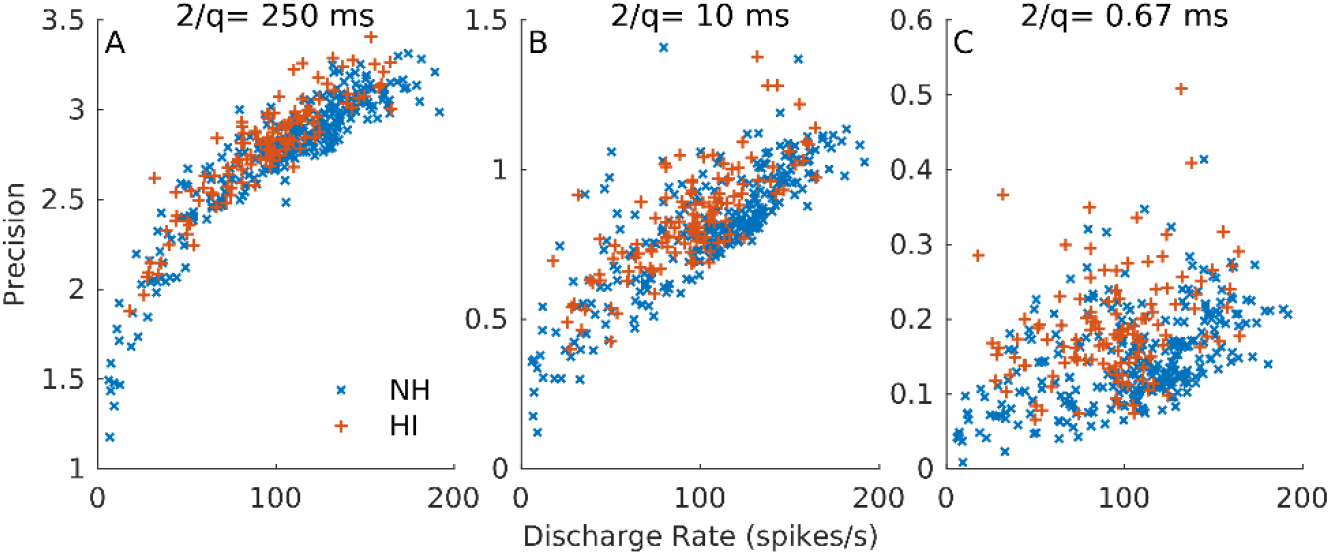
Temporal-coding precision for natural speech was not degraded by NIHL. (A) Across-trial precision is plotted versus discharge rate in response to a natural speech sentence in quiet at conversational levels. A time-shift cost (*q*) corresponding to 250-ms time scale was used to represent syllabic rate. Symbols represent individual AN fibers across all CFs. (B) 10-ms time scale to emphasize voice-pitch coding. (C) 0.67-ms time scale to emphasize speech-formant coding. A small but significant *increase* in temporal precision is observed for HI responses (group, F=30.8, p=5.4×10^−8^), which likely derives from increased responses to lower-frequency energy due to distorted tonotopy (CF×group interaction, F=5.0, p=0.025).

### Changes in AN-fiber tuning following NIHL distort the normal tonotopic representation of natural vowels

Normal AN fibers are characterized by high sensitivity (low threshold) and spectral specificity (sharp tuning), which allows for selective responses to stimulus energy near their CF. These NH properties produce tonotopic responses to complex sounds, as demonstrated previously for synthetic steady-state vowels (Miller et al., 1997; Young and Sachs, 1979). Here, we evaluated the effects of NIHL on natural-vowel responses (Figures 3 and 4) by examining spectral estimates [*D*(*f*)] computed from difference PSTHs (which minimize rectifier-distortion artifacts; Parida et al., 2021). The exemplar NH AN-fiber response shown in Figure 3 was dominated by the second formant F_2_, the spectral feature closest to fiber CF for the segment considered (Figure 3B). This effect can be seen in the NH *D*(*f*) (Figure 3C), which shows a clear peak near F_2_ (blue arrow), representative of a tonotopic response. In contrast, the exemplar HI fiber with similar CF had a *D*(*f*) with a peak near F_1_ (~ 650 Hz), which is well below its CF (~1.5 kHz, Figures 3B and 3C). This HI nontonotopic response could arise from some combination of its elevated tip threshold, broadened tuning, and hypersensitive tail, which are typical for AN fibers following NIHL (Liberman, 1984; Liberman and Dodds, 1984a). These physiological changes lead to a reduced TTR in the FTC (double arrows in Figure 3B), which is a consistent trend in our HI population (Figure 3D).

**Figure 3.**
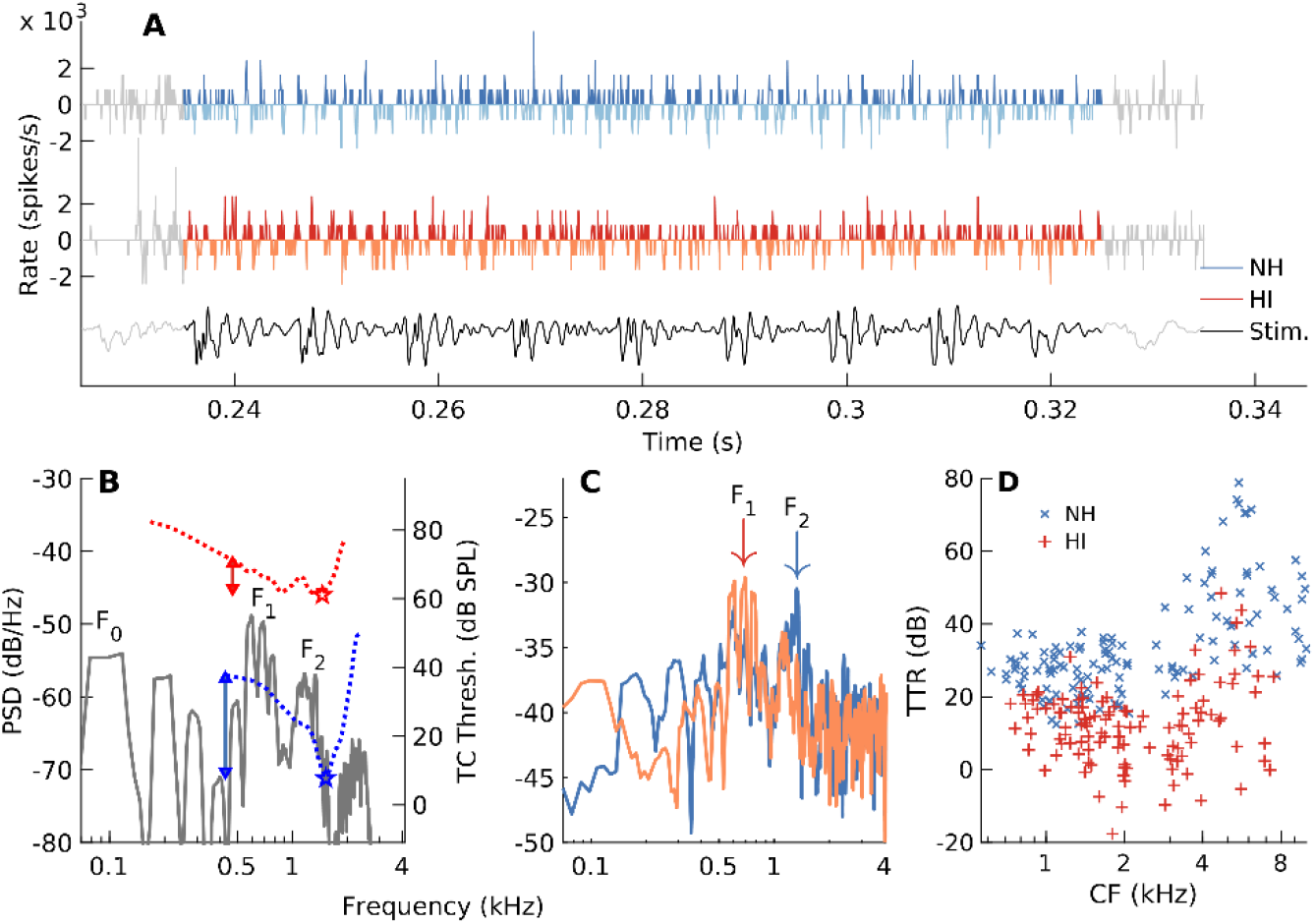
Changes in AN-fiber tuning following NIHL distort the normal tonotopic representation of natural vowels. (A) Alternating-polarity PSTHs in response to a “quasi-stationary” segment from a natural sentence in quiet (stimulus: dark gray). Darker (lighter) color shades represent PSTHs for positive (negative, reflected across x-axis for display) stimulus polarity. Stimulus scaled arbitrarily. (B) Spectrum of the stimulus segment (gray, left y-axis) with fundamental frequency (F_0_) and first and second formants (F_1_ and F_2_) labeled. Tuning curves (right y-axis) from the two example AN fibers in A and C. Stars indicate CF (where the HI CF was chosen based on convention from anatomical labeling studies; Liberman and Dodds, 1984c); double-arrows indicate tip-to-tail ratio (TTR) in dB. Both AN fibers have CF near F2. (C) Spectra of the difference PSTHs in A [*D*(*f*), representing NH and HI TFS responses]. NH *D*(*f*) shows a tonotopic response (i.e., near CF), with a peak near F2 (blue arrow); HI *D*(*f*) has a peak near F1 (red arrow), despite having CF near F2 (i.e., nontonotopic response). (D) TTR distributions for NH and HI fibers. TTR was consistently (group×CF interaction, F=2.9, p=0.09) and significantly (group, F=59.1, p=3.4×10^−6^) reduced following NIHL.

**Figure 4.**
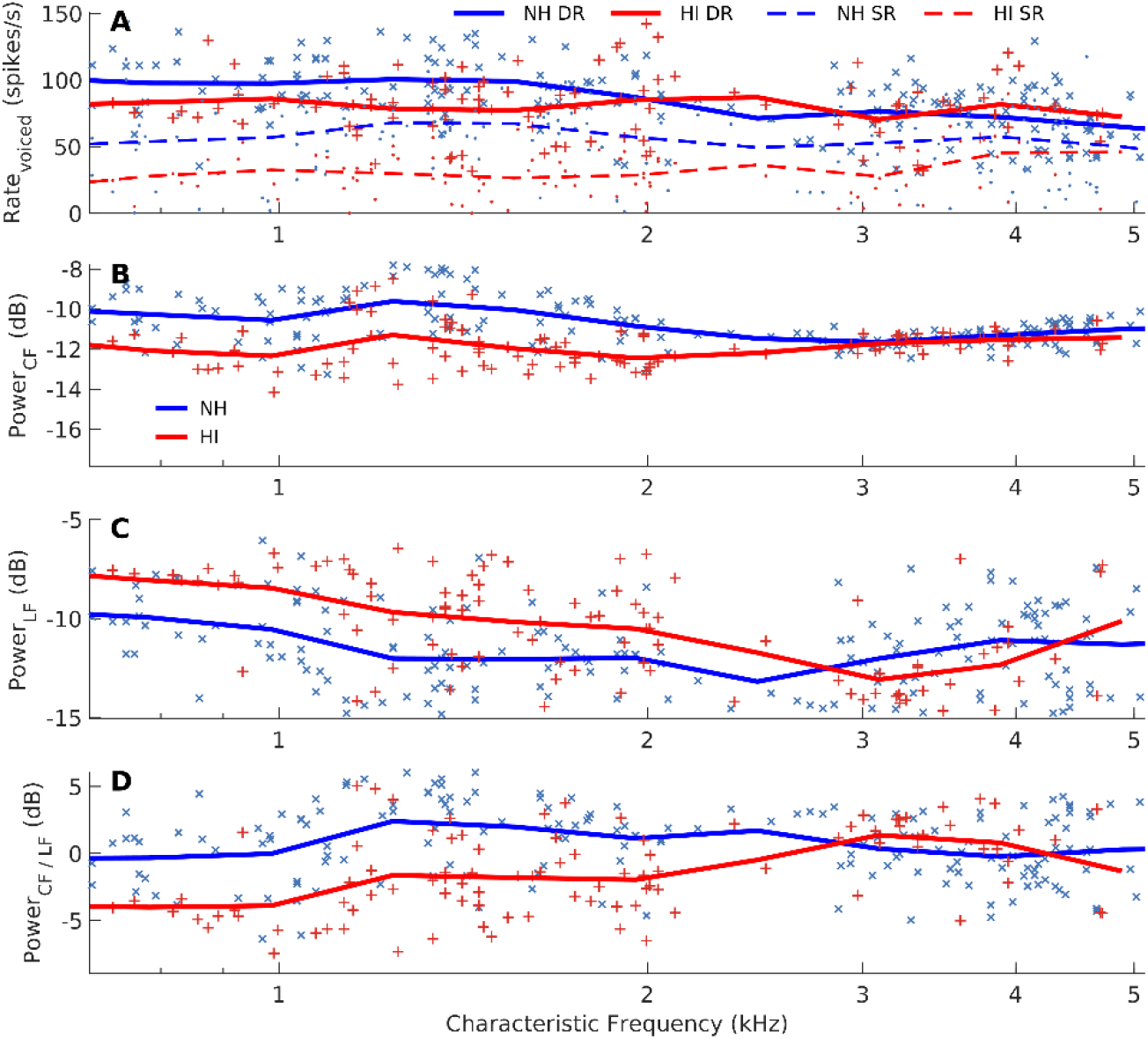
Distorted tonotopy enhances below-CF coding of voiced speech at the expense of near-CF representations, even when audibility is restored. (A) Driven rates (DR, solid trend lines) for NH and HI AN fibers were comparable in voiced portions of the natural speech sentence in quiet (group, F=0.6, p=0.44). Spontaneous rates (SR, dashed lines) were reduced for HI (F=44.1, p=1.5×10^−10^). Triangular-weighted trend lines (here and in the following figures) were computed for 2/3-octave averages. (B) Fractional response power near CF [in dB re: total power based on the difference-PSTH spectrum, *D*(*f*)] was significantly reduced for HI primarily at lower (e.g., <3 kHz) CFs (group, F=84.14, p<2×10^−16^; group×CF interaction, F=13.8, p=2.4×10^−4^). (C) Fractional response power in a LF band (<400 Hz) was enhanced for HI (group, F=10.5, p=1.3×10^−3^; group×CF interaction, F=23.0, p=2.5×10^−6^). (D) Ratio of power near CF to power in the low-frequency band was significantly reduced for HI fibers (group, F=36.86, p=3.9×10^−9^; group×CF interaction, F=26.7, p=4.5×10^−7^). CF range: 0.6-5 kHz in all panels.

### Distorted tonotopy enhances below-CF coding of voiced speech at the expense of near-CF representations, even when audibility is restored

To quantify the effects of distorted tonotopy on voiced-speech coding at the population level, fractional power near-CF and at low frequency (LF) were quantified as metrics related to tonotopic coding (Figure 4). Fractional power was used to minimize the effects of overall power and rate differences, if any. Audibility was not a primary contributing factor as driven rates during voiced segments were similar between the two groups (Figure 4A), despite the expected reduction in spontaneous rate (SR, Liberman and Dodds, 1984b).

Near-CF power was significantly lower for HI fibers, particularly at lower (< 3 kHz) CFs. To quantify the susceptibility of fibers with higher CFs (> 0.6 kHz) to very LF stimulus energy, the power in *D*(*f*) was computed in a low-pass spectral window (400-Hz 3-dB cutoff). As expected, LF power was significantly higher for HI fibers than for NH fibers (Figure 4C). The ratio of power near CF to power at LF, which indicates the strength of tonotopic coding (relative to LF coding), was > 0 dB on average for NH fibers with CF ≤ 3 kHz. In contrast, this relative-power metric was < 0 dB on average for HI fibers in the same CF region. Overall, these results demonstrate a disruption of near-CF stimulus energy coding at the expense of very LF stimulus energy coding at a single AN-fiber level. These deficits are more severe than previously thought (Henry et al., 2016), likely because of the steep negatively sloping spectrum of natural speech.

What physiological factors contribute to this distortion in tonotopic coding? To address this question, a mechanistic linear mixed model was constructed for relative near-CF to LF power (Figure 4D) as the response variable, with threshold, local Q_10_, and TTR included as predictors (in addition to the significant dependence on CF, F=11.7, p=7.8×10^−4^). Whereas traditionally hypothesized factors underlying speech-coding deficits include elevated threshold and broader bandwidth, the model suggested that threshold was a significant factor (d_eq_=0.35, p=.01), but Q_10_ was not (p=0.46). Interestingly, TTR was the major factor underlying this distorted tonotopic coding (d_eq_=0.42, p=3×10^−3^).

### Temporal-place formant representation was more susceptible to background noise following NIHL

The effects of distorted tonotopy on formant coding were evaluated for natural speech in quiet and in noise using harmonicgram analyses (Parida et al., 2021). Coding strength for the first three formants was quantified as the *D*(*f*)-based power estimate along the dynamic formant trajectories at different SNRs (Figure 5). For NH speech-in-quiet responses, fractional power for individual formant trajectories peaked at CFs near or slightly above the formants’ mean frequencies (Figure 5A, 5D, and 5G), a result consistent with previous reports of tonotopic formant coding (Delgutte and Kiang, 1984a; Young and Sachs, 1979). As expected, F_1_ representation was expanded for the HI group extending to CFs substantially above F_1_, (i.e., CFs in the F_2_ and F_3_ ranges). Coding of F2 and F3 was diminished for the HI group, and the corresponding peaks were shifted to higher CFs than expected based on the formant frequencies. This distorted tonotopy for natural speech is consistent with system characterization studies that report a lowering of best frequency following NIHL (Henry et al., 2016; Liberman, 1984), and with previous reports using synthesized vowels (Miller et al., 1997).

**Figure 5.**
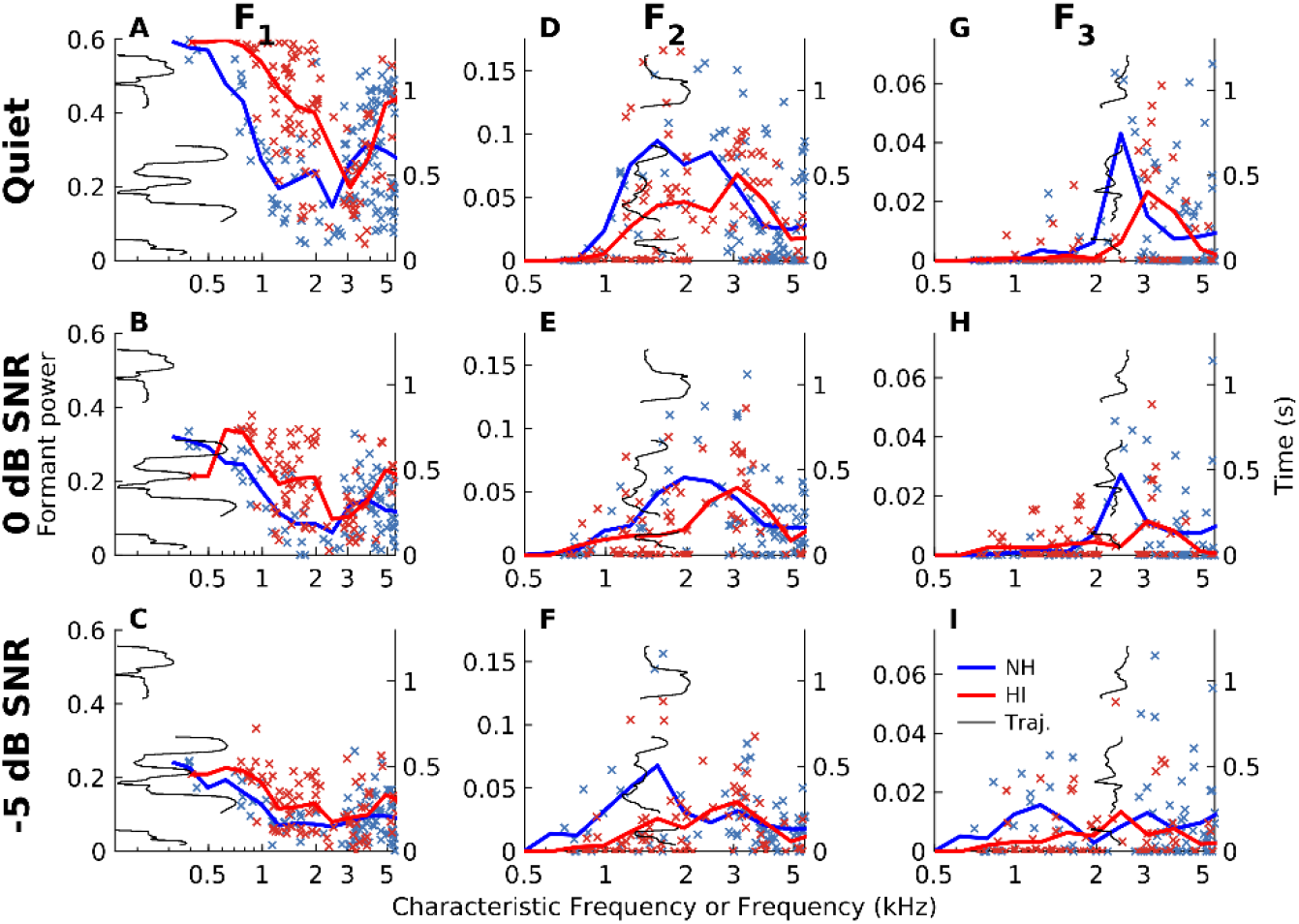
Temporal-place formant representation was more susceptible to background noise following NIHL. (A, B, and C) Temporal-coding strength for the first formant (F_1_) is shown for NH (blue) and HI (red) AN-fiber responses to the natural speech sentence in quiet (A) and in noise at SNRs of 0 and −5 dB (B, C). Markers represent harmonicgram-based fractional power estimates (left y-axis) along the corresponding formant trajectory (thin black lines, right y-axis) for the speech response relative to the fractional power for the noise-alone response. F1 coding was significantly enhanced for HI fibers (F=25.8, p=5.4×10^−7^; CF range: 0.5-5 kHz). (D, E, and F) Same as (A-C) but for F2. For the NH population, trend lines peak near formants, as expected for tonotopic coding. In contrast, HI trend lines peaked at frequencies well above F2. Compared to NH, F2 coding was degraded for HI (group, F=23.4, p=2.7×10^−6^; CF range: 1-2.5 kHz). (G, H, and I) Same layout for F_3_ coding, which was significantly degraded for HI (F=7.3, p=0.01; CF range: 2-3 kHz).

Noise had a strong detrimental effect on the already degraded (reduced strength and nontonotopic) representations of F_2_ and F_3_ for the HI population. At 0dB SNR, F_2_ and F_3_ peaks were still strong and tonotopic for the NH pool (Figure 5E and 5H). For HI fibers, the F_2_ peak was still discernible and centered at a CF location much higher than F_2_, but F_3_ coding was almost nonexistent. Similarly, at −5 dB SNR, the NH pool still showed a discernible and tonotopic peak for F_2_ coding, whereas F_2_ coding for the HI pool was severely degraded.

To identify the contribution of various physiological factors to these formant-coding degradations, mixed models were constructed for each formant fractional power response with TTR, Q10, and threshold as fixed-effect predictors, in addition to log-CF and SNR. For F_1_ enhancement in the HI representation TTR (d_eq_=0.21, p=0.034) was the major contributing factor and not threshold (d_eq_=0.19, p=0.051) or Q_10_ (p=0.26). Similarly for the degraded F_2_ response TTR (d_eq_=0.78, p=2.1×10^−6^), but not threshold (d_eq_=0.33, p=0.053) or Q_10_ (p=0.17), was the major factor. For F_3_, TTR (d_eq_=0.80, p=0.022) was the only contributing factor to the degraded response (threshold: p=.20; Q_10_: p=0.71).

Overall, these results show a degradation of higher-formant (F_2_ and F_3_) coding, which is known to be important for the perception of vowels as well as consonants, in HI-fiber responses following acoustic insult. Moreover, these degradations are more susceptible to the presence of noise, which is consistent with the stronger perceptual deficits that listeners with SNHL experience in noisy environments.

### Unlike voiced segments, driven rates for the fricative /s/ were not restored in HI fibers despite compensating for overall audibility loss

Fricatives constitute a substantial portion of phoneme confusions among listeners with SNHL (Bilger and Wang, 1976; Dubno et al., 1982; Van de Grift Turek et al., 1980). To explore potential neural bases underlying these deficits, neural responses to a fricative (/s/) were analyzed. Previous studies have reported robust fricative coding by NH AN fibers in terms of onset and sustained responses (Delgutte and Kiang, 1984b). NH fibers with higher CFs (i.e., near frequencies where /s/ has strong energy, Figure 6A) showed a sharp onset, followed by a robust sustained discharge (e.g., Figure 6B). In contrast, HI fibers showed a substantial reduction in onset response (e.g., Figure 6B), with less of an effect on the sustained response. This reduction in onset response contrasts with previous studies that reported increased onset responses to tones following NIHL; however, those results were for equal sensation level (Scheidt et al., 2010).

**Figure 6.**
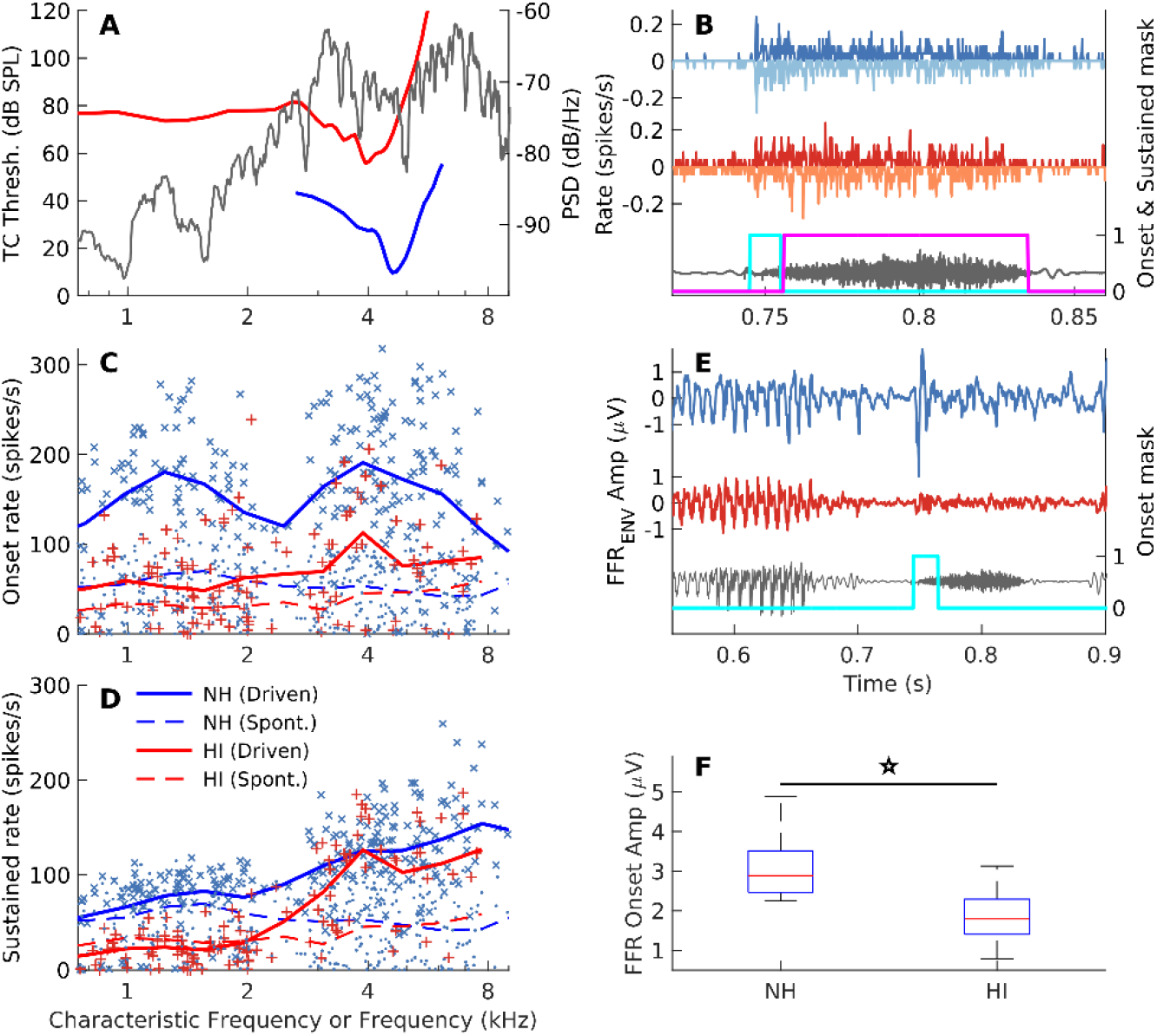
Unlike voiced segments, driven rates for the fricative /s/ were not restored in HI fibers despite compensating for overall audibility loss. (A) Spectrum of /s/ (gray) from the natural sentence and exemplar tuning curves. (B) Time-domain waveforms of /s/ (gray) and alternating-polarity PSTHs of AN fibers shown in A. Same format and data-analysis parameters as Fig 3A. Cyan (10-ms long) and magenta (80-ms) temporal windows denote the masks used to compute onset and sustained response rates. While these example AN fibers had comparable sustained rates, the HI onset response was substantially degraded. (C) Driven onset rates for the HI population were significantly reduced compared to NH for all CFs. Same format as Fig 4A (group, F=120.8, p<2×10^−16^; group×CF interaction, F=2.0, p=0.15). (D) Driven sustained rates were also reduced for the HI group in the high-CF (>2.5 kHz) region where /s/ had substantial energy (F=7.0, p=0.009). (E) Exemplar FFR_ENV_ data from a NH and a HI chinchilla demonstrate a reduced onset response for /s/ following NIHL. Cyan window (20-ms) indicates the onset mask. (F) Distributions of peak-to-peak onset amplitudes show a significant reduction in FFR onset response for HI chinchillas (F=5.8, p=0.039).

Driven onset rate was significantly lower for the HI population at all CFs (Figure 6C), whereas sustained rate was only slightly, but significantly, reduced for the HI group (Figure 6D).

To evaluate the noninvasive correlates of these single-fiber degradations in consonant coding, frequency-following responses (FFRs) were recorded in response to the speech stimulus from the same set of animals. Representative FFR responses from two animals (one from each group) reiterate these fricative onset-response degradations (~0.75 s in Figure 6E). While the NH FFR had a sharp onset, the HI FFR lacked any clear onset response. In contrast, responses during voiced speech were comparable between the NH and HI examples (e.g., ~0.6 s), mirroring the AN result that driven rates to voiced speech were similar between the two groups (Figure 4A). Thus, despite audibility restoration for voiced speech, there was a significant reduction in fricative onset response in the HI group (Figure 6F).

### Driven rates for stop consonants (/d/ and /g/) were also not restored despite overall audibility compensation

Stop consonants are among the most confused phonemes for listeners with SNHL (Bilger and Wang, 1976; Van de Grift Turek et al., 1980). The neural representations of two stop consonants (/d/ and /g/) present in the speech stimulus (Figure 7A) were also evaluated based on onset and sustained rates. In response to /d/ and /g/, the NH AN fiber showed a strong onset response, followed by sustained activity that was well-above spontaneous activity (Figures 7B and 7C). In contrast, for the HI AN fiber, both onset and sustained activity were substantially reduced. Population results show that onset and sustained rates for /d/ and /g/ were significantly degraded for the HI population relative to NH (Figures 7D–7G).

**Figure 7.**
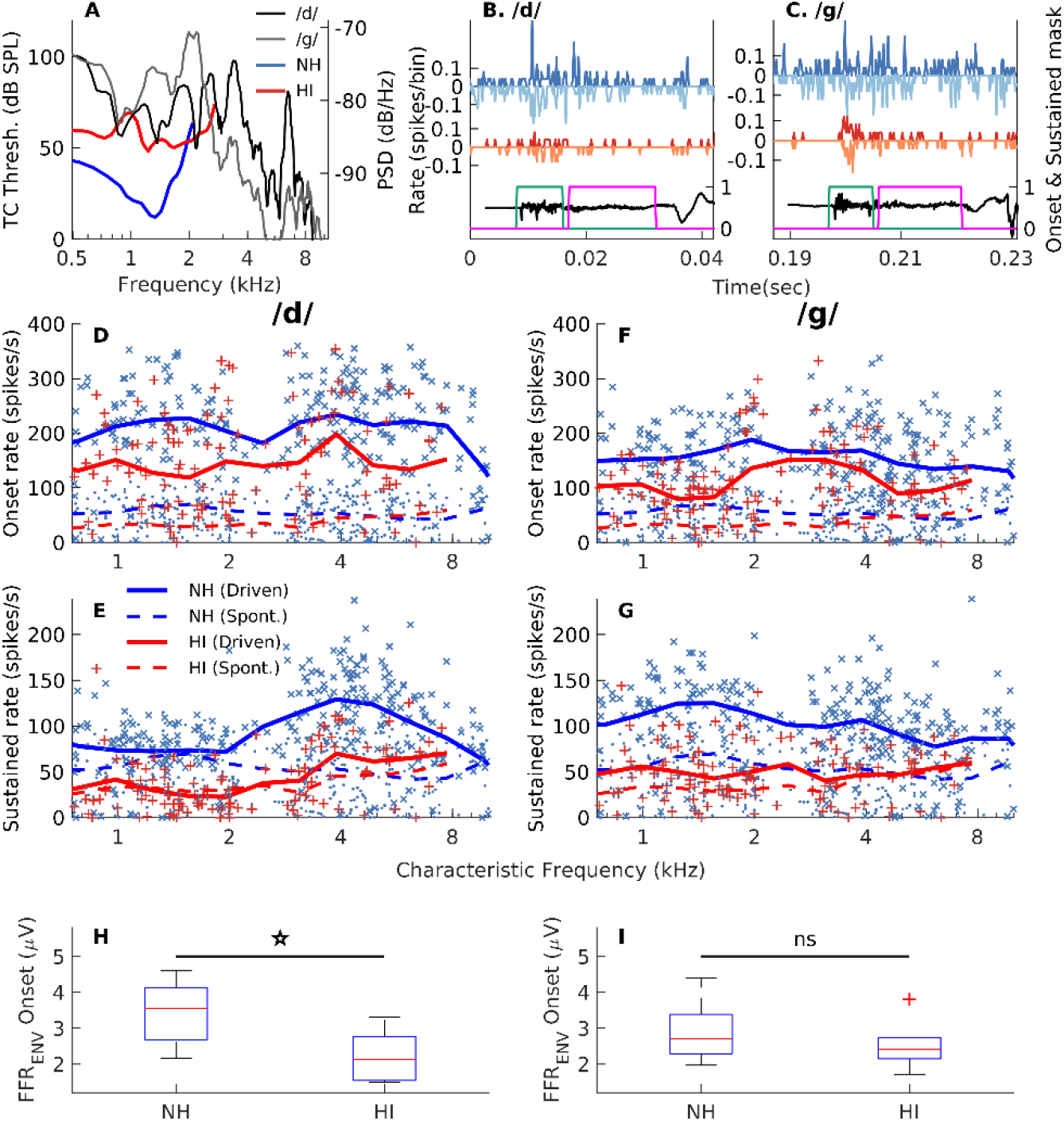
Driven rates for low-intensity stop consonants (/d/ and /g/) were also not restored despite overall audibility compensation. (A) Spectra for stop consonants /d/ (dark gray) and /g/ (light gray) from the natural sentence in quiet. FTCs of representative NH and HI AN fibers. (B) Alternating-polarity PSTHs for the two fibers in A in response to /d/. Same format and analysis parameters as Fig 6B. Both onset and sustained rates are reduced for the HI fiber. (C) Same layout but for /g/, which shows the same general effects as /d/. The onset/sustained windows in (B) and (C) were 8-/15-ms long. (D) Onset rates in response to /d/ were significantly reduced in the HI population compared to NH (F=57.9, p=2.3×10^−13^). Same format as Fig 6C. (E) Same layout and results as D but for /d/ sustained rates (F=149.8, p<2.2×10^−16^). (F and G) Same layouts and results as D and E but for /g/. Onset (F=41, p=1.1×10^−9^) and sustained (F=132.4, p<2×10^−16^) rates for /g/ were also reduced for the HI group. (H) Distributions of FFR peak-to-peak onset amplitudes in response to /d/ for chinchillas in both groups show a significant reduction for the HI group (F=5.97, p=0.037). (I) Same layout as H but for /g/. The observed reduction in FFR onset amplitudes was not significant (F=0.58, p=0.47). CF range for statistical analyses: 0.5-8 kHz for /d/ and 0.5-3 kHz for /g/.

FFR onset was significantly reduced for /d/ following NIHL (Figure 7H), consistent with the universal reduction in onset rate across the whole CF range for HI fibers (Figure 7D). The FFR onset response to /g/, however, was only slightly reduced (not significantly) for the HI group (Figure 7I). Overall, these FFR data (and those from Figure 6) suggest that degraded consonant representations, even when vowel audibility is restored, persist despite central-gain related changes that can occur in the midbrain (Auerbach et al., 2014).

### Changes in tuning following NIHL eliminate the noise-resilient benefits of AN fibers with lower SR for fricative coding

As previously described, listeners with SNHL often struggle in noisy environments in identifying consonants more so than vowels. Here, we investigated the effect of background noise on coding of the fricative /s/, which elicited robust sustained activity for both groups in the quiet condition (Figure 6D). When the sentence is mixed with noise at a particular SNR, even negative SNRs, the resultant signal often has specific spectrotemporal regions with a favorable SNR (e.g., high-frequency region for /s/, Figure 8A). These high-SNR regions likely mediate robust speech perception in noise (Cooke, 2006). In our data, NH AN fibers that were narrowly tuned near this high-SNR region responded selectively to the fricative energy (e.g., Figure 8B). In contrast, HI AN fibers showed reduced TTR (Figures 8A and 3D). As a result, HI fibers tuned to higher frequencies responded poorly to fricative energy and strongly to LF energy in either the speech (e.g., voiced segments) or noise (e.g., Figure 8C).

**Figure 8.**
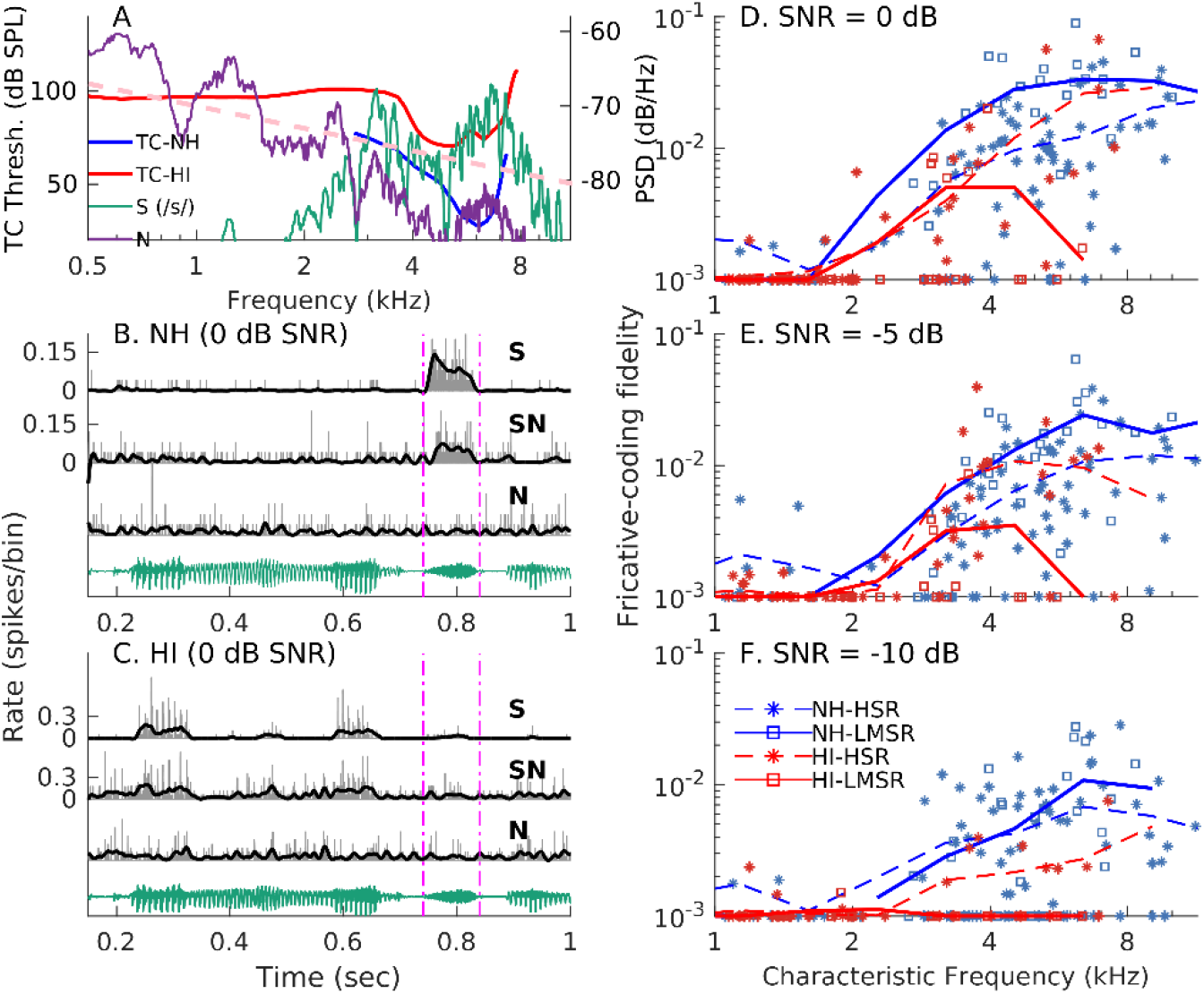
Changes in tuning following NIHL eliminate the noise-resilient benefits of AN fibers with lower SR for fricative coding. (A) Spectra for /s/ from the natural-speech sentence and the concurrent speech-shaped noise (N) segment for 0 dB overall SNR. Although overall SNR was 0 dB, a local high-SNR (10-15 dB) region occurs at high frequencies (> 3 kHz). FTCs of two example AN fibers that schematize the increased deleterious effect of speech-shaped background noise following NIHL, especially compared to an ideal pink noise (pink line). (B) PSTHs (grey) in response to speech-alone (S), noisy-speech (SN), and noise-alone for the NH fiber (SR=0.2/s) in A. Thick black curves represent response envelopes. Dashed pink lines indicate temporal region in the stimulus (green) containing /s/. (C) Same layout as B but for the HI fiber (SR=1.1/s). (D, E, and F) Speech-in-noise coding fidelity for /s/ at perceptually important SNRs, as quantified by the corrected correlation between responses to S and SN (minimum value set to 0.001 for display). Squares and asterisks correspond to AN fibers with low/medium SR (<18 /s) and high SR (>18 /s), respectively. For NH, AN fibers with low/medium SR show better coding in noise than high-SR fibers; however, the opposite is true following NIHL because the noise resilience of low/medium SR fibers was lost resulting in overall degraded fricative fidelity (SR, F=16.5, p=6.2×10^−5^; group, F=3.3, p=0.07; group×SR interaction, F=18.8, p=2.0×10^−5^).

Fricative-coding fidelity was quantified by the *corr_corrected_* metric, defined as the difference between the response-envelope correlations for speech-alone and noisy-speech, and speech-alone and noise-alone. Responses of AN fibers with low/medium SR, hypothesized to drive robust human speech perception in noise (Bharadwaj et al., 2014), were particularly resistant to noise in the NH group compared to high-SR fibers (Figure 8D and 8E). In contrast, coding fidelity for low/medium-SR fibers was significantly lower than high-SR fibers following NIHL.

To identify the contribution of various physiological factors to fricative coding in noise, a mixed model was constructed with *corr_corrected_* as the response variable and with TTR, Q10, and threshold as fixed-effect predictors, in addition to log-CF and SNR. Results indicated that TTR (d_eq_=0.34, p=0.028) had a greater effect compared to Q_10_ (d_eq_=0.29, p=0.053), whereas threshold did not contribute significantly (p=0.54).

## DISCUSSION

A common statement from patients with SNHL after receiving a hearing aid is “I’m so grateful I can hear you now, but I still can’t fully understand you, especially in background noise.” Several neurophysiological mechanisms are hypothesized to underlie these suprathreshold deficits; however, neurophysiological studies examining the effects of SNHL on speech coding have been typically limited to synthesized speech tokens without background noise. Here, we describe the first data characterizing the effects of SNHL on AN-fiber responses to a natural speech sentence in the presence of background noise. These data elucidate physiological mechanisms that do and do not contribute to deficits in peripheral coding of natural speech following SNHL. In particular, these data highlight the prominent role that distorted tonotopy plays in degrading the coding and increasing the noise susceptibility of both vowels and consonants.

### Commonly hypothesized suprathreshold deficits were not the major factors degrading the neural coding of natural speech following NIHL

The two suprathreshold deficits most commonly hypothesized to affect speech perception in noise are degraded frequency selectivity (Glasberg and Moore, 1989; Halliday et al., 2019; Horst, 1987) and diminished temporal-coding precision (e.g., Hopkins et al., 2008; Lorenzi et al., 2006; Strelcyk and Dau, 2009); however, these hypotheses have not been evaluated in peripheral neural responses to natural speech following SNHL. Furthermore, previous neurophysiological studies of the effects of SNHL on vowel coding often confounded broadened tuning at the FTC tip and hypersensitive tails by quantifying the sharpness measure Q_10_ based on the broadest bandwidth 10 dB above threshold, i.e., including both tip and tail effects in one metric (Henry and Heinz, 2012; Henry et al., 2016; Kale and Heinz, 2010; Miller et al., 1997).

Here, we separated these effects by quantifying Q_10_ locally to represent only the broadening of the FTC tip, more consistent with the classic definition of frequency selectivity based on the auditory filter at CF. We found that this local Q_10_ was not a significant factor in degraded voiced-speech coding in quiet (Figure 4). This result is consistent with perceptual studies suggesting that degraded frequency selectivity may not be the primary factor explaining degraded speech perception in quiet (Dubno and Dirks, 1989; Festen and Plomp, 1983). More surprisingly, we found that TTR and not Q_10_ was the most significant factor in explaining the increased susceptibility to noise of formant (Figure 5) and fricative coding (Figure 8).

We also found that there was no degradation in the temporal-coding precision of AN-fiber responses to natural speech (Figure 2). Although some studies have reported degraded temporal coding (phase locking) after SNHL (Woolf et al., 1981), our result is consistent with the majority of studies that have not found degraded phase locking to pure tones in quiet following SNHL (e.g., Harrison and Evans, 1979; Henry and Heinz, 2012; Miller et al., 1997). Our quantification of temporal precision based on the average across-trial Victor-Purpura distance (Victor and Purpura, 1996) in Figure 2 included all CFs from our fiber population, and as such would have been sensitive to any shift in the underlying roll off of phase locking versus CF (e.g., reduced roll-off frequency, or reduced maximum phase-locking strength). Thus, these natural-speech coding data, along with the majority of previous studies using laboratory stimuli, provide no evidence for a reduction in the fundamental ability of AN fibers to encode rapid acoustic fluctuations.

That being said, changes in phase-locking strength in response to complex sounds can be observed following SNHL [e.g., a decrease in the balance of fine-structure to envelope coding resulting from enhanced envelope coding (Kale and Heinz, 2010); a decrease in tone phase locking in background noise due to broadened tuning (Henry and Heinz, 2012); or a small but significant *increase* in temporal precision (Figure 2) resulting from increased coding of lower-frequency energy due to distorted tonotopy]. Although these changes in temporal-coding strength of complex signals may be perceptually relevant, these temporal-coding effects occur due to factors other than a change in the fundamental ability of AN fibers to respond to rapid fluctuations.

### Distorted tonotopy was the major factor in degraded coding and increased noise susceptibility for both vowels and consonants following NIHL

The primary factor affecting degraded coding of natural speech in noise was the loss of tonotopy, a hallmark of normal cochlear processing. This degradation in the coding of complex sounds is associated with the loss of FTC tips and/or the hypersensitivity of FTC tails following NIHL (Figure 3). For voiced-speech segments where the driven rates in HI AN fibers were comparable between the two groups, the spectral content in the responses differed substantially in terms of tonotopicity. Our results showed that TTR was the only dominant physiological factor explaining the increased representation of LF energy at the expense of near-CF energy for voiced segments of speech (Figure 4). Our analysis of this effect on formant coding in dynamic natural speech required a significant advance in neural analyses using frequency demodulation applied to alternating-polarity PSTHs (i.e., the harmonicgram, Parida et al., 2021), because standard Fourier analysis blurs the coding of dynamic formants. These results at the single-fiber level are also consistent with our recent FFRs to natural speech in NH and HI chinchillas, which showed that very LF energy (<250 Hz) was overrepresented in the more central evoked responses (Parida and Heinz, 2021). In fact, those analyses show that the degree of distorted tonotopy is greater in portions of speech that have more negatively sloping spectra.

Another significant and perceptually relevant effect of distorted tonotopy was the increased susceptibility to background noise following NIHL. Increased noise susceptibility in HI fibers was seen for both the spectral coding of voiced speech (Figure 5) as well as for fricative coding (Figure 8). Our data on fricative coding in speech-shaped noise provide unique insight into the real-world significance of this effect. The fricative /s/, which has primarily high-frequency energy, normally has a better-than-average SNR due to the steep spectral decay of speech-shaped noise and the tonotopic coding in the NH cochlea. Without this normal tonotopicity following SNHL, the SNR in AN fibers with CFs within the spectral band of the fricative is greatly reduced due to the substantial LF noise now driving the neural response. Based on our previous FFR results illustrating the dependence of distorted tonotopy degree on spectral timbre (Parida and Heinz, 2021), this deleterious effect on fricative coding is expected to be greater for speech-shaped noise than for white or even pink background noise. While many environmental sounds have more LF energy consistent with a roughly pink spectrum (−3 dB/octave drop in energy), long-term speech has a downward spectral slope that is roughly 3 times steeper than pink noise (Figure 8A; also see French and Steinberg, 1947). Thus, these neural results have important implications for listening in the presence of multiple talkers, which is a condition of great difficulty for many listeners with SNHL (Festen and Plomp, 1990; Le Prell and Clavier, 2017).

### Noise-resistant ability of low/medium SR fibers for fricative coding was substantially degraded following NIHL

Beyond the underlying mechanisms for the increased noise-susceptibility of speech coding following NIHL, our data are insightful for fundamental speech coding in noise and have implications for other forms of hearing loss. The statistical analyses of our fricative-in-noise coding data (Figure 8) showed a significant effect of SR, and an interaction between group and SR. The main effect of SR was not surprising given previous reports from NH studies of the superior coding-in-noise of low-SR fibers (Costalupes et al., 1984; Young and Barta, 1986), including for consonant-vowel tokens (Silkes and Geisler, 1991). The new insight provided by our data relates to the interaction between group and SR, which arises because NIHL caused a larger deficit in speech fidelity in the low/medium-SR fibers compared to the normally poorer high-SR fibers. These results are particularly important as recent studies on the effects of age and noise exposure on cochlear synaptopathy show that older listeners with or without noise exposure are likely to have fewer low/medium-SR fibers remaining (Fernandez et al., 2015, 2020; Wu et al., 2019).

### Audibility restoration is not the same for consonants and vowels due to expanded AN-fiber threshold distribution following NIHL

Psychoacoustic studies suggest that reduced fricative perception for listeners with SNHL depends on both reduced ability to use formant transitions and a reduced audibility of the frication cue (Zeng and Turner, 1990). As discussed for vowels, the dynamic representations of transitions for higher formants (F2 and F3) were substantially degraded for the HI population, especially in noise (Figure 5). Formant transitions are also important for speech-in-noise perception of low-intensity stop consonants that are easily masked by noise; thus, the increased noise susceptibility of these transitions likely contributes to these perceptual deficits.

The audibility of consonants in our study, as indexed by driven onset and sustained rates (Figures 6 and 7), was often not restored, in contrast to vowels (Figure 4A). Such divergent audiometric effects are consistent with psychoacoustic studies that report similar audibility differences for consonants and vowels even after compensating for overall audibility (Phatak and Grant, 2014). This reduced audibility following NIHL likely contributes to consonant confusions, which are common for both fricatives and stop consonants (Bilger and Wang, 1976; Turner and Robb, 1987; Zeng and Turner, 1990).

Our data suggest that the expanded AN-fiber threshold distribution following NIHL (Figures 1E and 1F) may contribute to differences in audibility of consonants and vowels in amplified speech. Although this finding is contrary to assumptions made in the psychophysical literature on loudness recruitment (Moore et al., 1985; Zeng and Turner, 1991), consistently expanded threshold distributions have been observed in both chinchillas and cats following NIHL (Figures 1E and 1F; Heinz et al., 2005). Thus, compensating for audiometric loss only guarantees the restoration of the most sensitive AN fibers. Because the elevation in the 10^th^ percentile thresholds underestimated the elevation in the 50^th^ percentile, lower-amplitude consonants activated a smaller percentage of the population following NIHL, and thus were not as “audible” within the population despite restoration of vowel audibility.

### Variations in the degree of distorted tonotopy across etiologies may contribute to individual differences in real-world speech perception

There is general consensus among researchers regarding the inadequacies of the audiogram to account for real-life perceptual deficits. Individual variability in speech perception, even among listeners with similar audiograms, likely stems from variations in the degree of suprathreshold deficits. Here, we have demonstrated that distorted tonotopy is a significant factor in the degraded coding and increased noise susceptibility of natural speech following NIHL. Because the degree of distorted tonotopy appears to vary across different etiologies (e.g., NIHL or age-related hearing loss) even for similar degrees of hearing loss (Henry et al., 2019), it is likely that this variation contributes to individual difference in speech-perception deficits. The development of noninvasive diagnostics to identify distorted tonotopy (e.g., Parida and Heinz, 2021) is critical for determining the extent and perceptual relevance of this important physiological mechanism affecting the neural coding of natural speech.

## STAR*METHODS

Detailed methods are provided in the online version of this paper and include the following:

- KEY RESOURCE TABLE
- RESOURCES AVAILABILITY

- Lead contact
- Materials availability
- Data and code availability
- EXPERIMENTAL MODEL AND SUBJECT DETAILS
- METHOD DETAILS

- Noise exposure and electrophysiological recordings
- Surgical preparation and neurophysiological recordings
- Stimuli
- QUANTIFICATION AND STATISTICAL ANALYSIS

- Spike-train temporal-precision analysis
- Spectral analyses of alternating-polarity PSTHs
- Quantifying formant coding strength using the harmonicgram
- Coding fidelity metric for fricatives in noise based on correlation analysis
- Statistical analyses

## ACKNOWLEDGMENTS

We thank Kenneth Henry, Alex Hustedt-Mai, and Hari Bharadwaj for their valuable feedback on an earlier version of this manuscript. We also thank Josh Alexander for stimulating discussions on human speech perception. This research was supported by an International Project Grant (G72) from Action on Hearing Loss (UK) and by NIH (R01-DC009838).

## AUTHOR CONTRIBUTIONS

Conceptualization, S.P. and M.G.H.; Methodology, S.P. and M.G.H.; Software, S.P.; Formal Analysis, S.P.; Investigation, S.P.; Data Curation, S.P. and M.G.H.; Writing – Original Draft, S.P.; Writing – Review & Editing, S.P. and M.G.H.; Visualization, S.P.; Funding Acquisition, M.G.H.

## DECLARATION OF INTERESTS

The authors declare no competing interests.

## STAR*METHODS

### KEY RESOURCES TABLE

| REAGENT or RESOURCE | SOURCE | IDENTIFIER |
| --- | --- | --- |
| Deposited data |  |  |
| Chinchilla data | This study | https://github.com/HeinzLabPurdue/D_T_SpINcoding-paper |
| Cat data | Heinz et al., 2005 | https://github.com/HeinzLabPurdue/D_T_SpINcoding-paper/tree/main/CATdata |
| Experimental models: organisms/strains |  |  |
| Chinchillas (chinchilla lanigera) | Ryerson Chinchilla Ranch | N/A |
| Software and algorithms |  |  |
| MATLAB 2018b | MathWorks | N/A |
| R4.0.4 | https://www.r-project.org/ | N/A |
| Victor-Purpura distance | Chicharro et al., 2011 | N/A |
| Harmonicgram analysis | Parida et al., 2021 | https://github.com/HeinzLabPurdue/D_T_SpINcoding-paper |
| Other |  |  |
| Enclosed subwoofer | Selenium 10PW3 | Selenium Components |
| DSP boards | Tucker-Davis Technologies<br>National Instruments | RP2, RX8, HB7, PA5<br>PCIe-6341, PCIe-6612 |
| Speakers | Etymotic Research | ER2 |
|  | Beyerdynamic | DT48 |
| Microphone | Etymotic Research | ER10b or ER7c |
| Analog amplifiers/filters | World Precision Instruments | ISO-80 |
|  | Dagan Corporation | 2400A |
|  | Krohn-Hite Corporation | 3550 |
| Spike time/amplitude window discriminator | BAK Electronics | DIS-1 |

### RESOURCES AVAILABILITY

#### Lead contact

Further information and requests for resources should be directed to and will be fulfilled by the

Lead Contact, Michael Heinz.

#### Materials availability

This study did not generate new unique reagents.

#### Data and code availability

The datasets and code supporting this study are available at https://github.com/HeinzLabPurdue/DT_SpINcoding-paper.

### EXPERIMENTAL MODEL AND SUBJECT DETAILS

All procedures described below followed PHS-issued guidelines and were approved by Purdue University Animal Care and Use Committee (Protocol No: 1111000123). Male chinchillas (<1-year old, weighing between 400 and 700 gm) were used in all experiments. Animals were socially housed in groups of two until they underwent any anesthetized procedure, after which they recovered in their own cage. All animal received daily environmental enrichment. Animal facility was maintained in a 12-hour light/12-hour dark cycle.

### METHOD DETAILS

#### Noise exposure and electrophysiological recordings

Detailed descriptions of the noise-exposure and electrophysiological-recording procedures are provided elsewhere (Parida and Heinz, 2021), with brief descriptions provided here. A single discrete noise exposure (116 dB SPL [C-weighted], 2-hour duration, octave-band noise centered at 500 Hz) using an enclosed subwoofer (Selenium 10PW3, Harman; placed ~30 cm above the animal’s head) was used to induce NIHL. Noise levels were calibrated at the entrance of the ear canal using a sound-level meter (886–2, Simpson, Elgin, IL, USA). Animals were allowed at least two weeks to recover following noise exposure before any recordings were made. Animals were anesthetized using xylazine (2 to 3 mg/kg, subcutaneous) and ketamine (30 to 40 mg/kg, intraperitoneal) for data recordings and noise exposure. The rectal temperature of all anesthetized animals was maintained at 37°C using a feedback-controlled heating pad (50-7053F, Harvard Apparatus). Atipamezole (0.4 to 0.5 mg/kg, intraperitoneal) was used to facilitate faster recovery from anesthesia following noninvasive experiments.

ABRs and FFRs were recorded using three subdermal needle electrodes in a vertical montage (vertex to mastoid, differential mode, common ground near animals’ nose; Henry et al., 2011; Zhong et al., 2014). ABRs (bandwidth=0.3–3 kHz) and FFRs (30 Hz – 1 kHz) were bandlimited using analog filters (ISO-80, World Precision Instruments, Sarasota, FL; 2400A, Dagan, Minneapolis, MN). DPOAEs were measure using an in-ear microphone (Etymotic ER-10B, Etymotic Research, Elk Grove Village, IL, USA), following in-ear calibration for each animal. Calibrated sound was presented using ER2 speakers (Etymotic Research) for electrophysiological recordings. Sound presentation and data recordings were controlled using a custom-integrated system of hardware (Tucker-Davis Technologies, Alachua, FL; National Instruments, Austin, TX) and software (MATLAB, The MathWorks, Natick, MA).

#### Surgical preparation and neurophysiological recordings

Detailed surgical-preparation and neurophysiological-recording procedures are described by Henry et al. (2016), and are only briefly described here. Anesthesia was induced with the same doses of xylazine/ketamine used for ABRs and was maintained with sodium pentobarbital (~7.5 mg/kg/hour, intraperitoneal). Animals were supplemented with lactated Ringer’s solution during experiments (~1 ml/hour), which typically lasted 18-24 hours. A posterior fossa approach was employed for the craniotomy in the right ear, following venting of the bulla with 30 cm of polyethylene tubing to maintain middle-ear pressure.

Spike trains were recorded from single AN fibers of anesthetized chinchillas using glass micropipettes (impedance between 10 and 50 MΩ). Recordings were amplified (2400A, Dagan) and filtered from 0.03 to 6 kHz (3550, Krohn-Hite). Isolated spikes were identified using a time–amplitude window discriminator (BAK Electronics, Mount Airy, MD, USA) and stored digitally with 10-μs resolution. Experiments were terminated if sudden shifts in FTC threshold and tuning were observed for two or more AN fibers, following which the animal was sacrificed with Euthasol (2 ml, intraperitoneal; Virbac AH, Inc., Fort Worth, TX). Single-unit data are from 15 NH (286 AN fibers) and 6 HI (119 AN fibers) chinchillas.

#### Stimuli

##### Screening (ABR and DPOAE) experiments

For ABRs, tone pips (5-ms duration, 1-ms on and off ramp) ranging from 0.5 to 8 kHz (octave spaced) were played at 0 dB SPL to 80 dB SPL in 10-dB steps. 500 repetitions of both positive and negative polarities were played for each intensity condition. ABR threshold was calculated based on a cross-correlation analysis (Henry et al., 2011). Another intermediate (odd multiple of 5 dB) step was used near preliminary ABR threshold estimate to fine-tune the final estimate. DPOAEs were measured for pairs of tones (f_1_, f_2_) presented simultaneously with f_2_/f_1_ =1.2 at 75 (f_1_) and 65 (f_2_) dB SPL.

##### Frequency-following-response (FFR) experiments

A naturally spoken speech sentence (list #3, sentence #1) from the Danish speech intelligibility test [CLUE, (Nielsen and Dau, 2009)] was used for FFR experiments. Intensity was set to 70 dB SPL for both groups. Both positive and negative polarities (500 repetitions/polarity) of the stimulus were used to allow estimation of envelope and temporal fine structure components from the FFR. Both envelope (FFR_ENV_ = average of FFRs to opposite polarities) and temporal fine structure (half the difference between FFRs to opposite polarities) amplitudes were comparable between the two groups (also see Parida and Heinz, 2021). This restoration in amplitude at 70 dB SPL likely reflects convergence in audibility at moderate intensities for mild-moderate hearing loss as well as central-gain related changes in the midbrain, which substantially contribute to the FFR (King et al., 2016).

FFR_ENV_ was used to analyze onset responses in Figures 6 and 7. Only the onset response was considered to evaluate consonant coding because evoked responses like the FFR require synchronous activity (e.g., the onset) across populations of fibers. As sustained responses to fricatives lack a clear temporal pattern, they are rather weakly represented in the FFR (Skoe and Kraus, 2010). Onset strength was quantified as the peak-to-peak FFR amplitude in an onset window (e.g., Figure 6E).

##### AN experiments

Monaural sound was delivered via a custom closed-field acoustic system. A dynamic speaker (DT-48, Beyerdynamic, Farmingdale, NY, USA) was connected to a hollow ear bar inserted into the right ear canal to deliver calibrated acoustic stimuli near the tympanic membrane. Calibrations of the acoustic system was done at the beginning of the experiment using a probe-tube microphone (ER-7C, Etymōtic, Elk Grove Village, IL, USA) that was placed within a few millimeters of the tympanic membrane.

Single AN fibers were isolated by advancing the electrode while playing broadband noise (20-30 dB re 20 μPa/√Hz; higher as needed for noise-exposed animals) as the search stimulus. Monopolar action potential waveform shape was used to confirm that recordings were from AN-fiber axons (as opposed to bipolar shapes exhibited by cell bodies in the cochlear nucleus). Prior to collecting spike-train data in response to speech and/or noise, all AN fibers were characterized as follows. An automated algorithm was used to estimate the FTC (Chintanpalli and Heinz, 2007). FTCs were smoothed by a 3-point triangular window before estimating parameters such as CF, threshold, Q_10_, and TTR. For HI fibers, CF was determined as the local minimum closest to the steep high-frequency-side FTC slope, which closely matches the basilar membrane frequency map following NIHL (Liberman, 1984; Miller et al., 1997). The threshold at CF was considered the threshold of the AN fiber. FTC 10-dB bandwidth was estimated as the minimum linearly interpolated bandwidth near the CF corresponding to an FTC threshold equal to 10 dB above the CF threshold. This approach of estimating Q_10_ differs from previous studies that used the maximum bandwidth criterion (instead of our minimum bandwidth criterion), and enables us to decouple tuning-broadening effects from tonotopic-distortion effects (Henry and Heinz, 2012; Henry et al., 2016; Kale and Heinz, 2010; Miller et al., 1997). Q_10_ was estimated as the ratio of CF to 10-dB FTC bandwidth. For a minority of HI AN fibers, the FTC threshold at the lowest or highest frequency measured was below the 10-dB bandwidth criterion (i.e., CF-threshold + 10 dB); in those cases, the 10-dB bandwidth was based on the lowest or highest frequency measured, respectively. This likely led to a slightly overestimated Q_10_ value for a few very-broad HI AN fibers. Tonotopic-distortion effects were quantified using TTR, which was estimated as the difference (in dB) between the threshold at CF (i.e., the tip) and the lowest threshold for frequencies at least 1.5 octaves below CF (i.e., the tail). Spontaneous rate (SR) was measured over a 30-s silence period for each AN fiber.

For AN experiments, the same speech sentence was used as for the FFRs. The overall intensity was set to 65 dB SPL for NH chinchillas and 80 dB SPL for HI chinchillas. The spectrally flat gain of 15 dB was roughly based on the half-gain rule (Lybarger, 1978), which has been used in AN studies of NIHL (Schilling et al., 1998), and was confirmed to restore driven rates for voiced portions of the sentence in preliminary experiments. Speech was also presented after mixing with frozen steady-state speech-shaped noise at three different perceptually relevant SNRs (−10, −5, and 0 dB). The speech-shaped noise was spectrally matched to 10 sentences spoken by the same speaker using autoregressive modelling. Both polarities of stimuli were presented in an interleaved manner (25 trials per polarity for most AN fibers).

#### QUANTIFICATION AND STATISTICAL ANALYSIS

##### Spike-train temporal-precision analysis

Temporal precision of spike trains was measured using the Victor-Purpura (VP) distance metric (Victor and Purpura, 1996), which quantifies the dissimilarity between two sets of spike trains. If a fiber responds precisely and consistently across different stimulus trials, the VP distance between the two spike trains will be small. The VP distance between two spike trains, *X* (total spikes = *N_X_*) and *Y* (*N*_Y_), is defined as the total cost involved in transforming *X* so that it matches *Y*. This transformation allows three operations: (1) addition of spikes, (2) deletion of spikes, and (3) shifting of spikes. The cost of adding or deleting a spike is set to 1 by convention. The cost of shifting a spike by Δ*t* seconds is proportional to a time-shifting cost parameter (*q*), which controls the temporal resolution of the analysis. This is because when /Δ*t*/ ≥ *2/q*, the cost of shifting the spike is greater than simply deleting the spike and adding a new spike. Note that for *q=0*, this analysis simplifies to a rate code where the cost is simply the difference in the number of spikes |*N_X_ - N_Y_* |. As *q* increases, the emphasis on temporal coding becomes greater as the temporal resolution improves. Specifically, all time shifts |*Δt*/ ≥ *2/q*, are maximally costly (cost=2) in this analysis, which can be interpreted as saying this analysis is most sensitive to frequencies less than *q/2*. Another way to think about this is for a temporal resolution (sampling rate) of *1/q*, the Nyquist rate of frequencies considered is *q/2* Hz. We have used this time/frequency relation to pick the specific *q* values considered here (Figure 2) with respect to the three main temporal resolutions of importance for speech (Rosen, 1992).

For a given temporal resolution *1/q*, the most optimal (smallest total cost) set of operations to transform *X* to *Y* is determined based on dynamic programming; VP distance between *X* and *Y* is computed as the total cost of all operations involved in this optimal transformation. For each pair of spike trains *X* and *Y*, the maximum possible cost (*VP_max_*) to transform *X* into *Y* depends on the number of spikes, specifically equal to the sum of the number of spikes in each spike train, *N_X_ + N_Y_* (corresponding to removing all spikes in *X* and adding in all spikes in *Y*). The minimum value of VP distance is *VP_min_*=| *N_X_ - N_Y_*|, corresponding to the condition where *q* = 0, and the only cost consists of equalizing the number of spikes. To minimize the basic dependence of VP distance on the number of spikes, we computed a normalized VP distance:

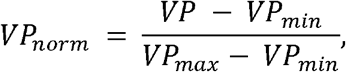

where, by definition, *VP_norm_* ranges from 0 to 1. Note that despite this normalization, some dependence on discharge rate remains because with more spikes in each spike train, it is more likely to find a nearby spike (i.e., the expected time shift between spikes is less when there are more spikes). As such, we compare VP distance (or precision) between groups as a function of discharge rate in Figure 2.

To compute the total VP distance for a given AN fiber and shifting cost parameter *q*, average *VP_norm_* was computed across all combinations of spike trains (i.e., across different stimulus trials). Temporal precision was then computed as ln(1/*VP_norm_*), where the natural logarithm was used to linearize the data for statistical analyses.

##### Spectral analyses of alternating-polarity PSTHs

Temporal and spectral analyses of spike-train responses were based on alternating-polarity PSTH analyses (Parida et al., 2021), with a PSTH bin width of 200 μs in all cases. To emphasize temporal-fine-structure responses for voiced speech coding in Figures 3–5, the difference PSTH, *d[n]*, was constructed by halving the difference between the PSTHs to opposite stimulus polarities. The difference-PSTH spectrum, *D*(*f*), was computed using multitaper (n=3) analyses in Figure 4 (Thomson, 1982). For each AN-fiber response, near-CF fractional power (in dB re: total response power) was quantified based on the *D*(*f*) power in a spectral window centered at CF. The spectral window was fourth order, and its 3-dB bandwidth was set to the 50^th^ percentile fit for AN-FTC 3-dB bandwidth for NH chinchillas (Kale and Heinz, 2010). 3-dB bandwidth was estimated from 10-dB bandwidth using the following formula, which provided a reasonable approximation for fourth order Butterworth filters.

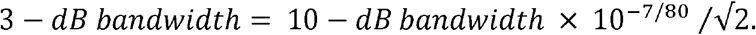

Low-frequency fractional power was quantified using a low-pass spectral window (cut-off=400 Hz, 10^th^ order).

##### Quantifying formant coding strength using the harmonicgram

Formant coding is traditionally quantified using the Fourier spectrum of the period histogram or the difference PSTH (Sinex and Geisler, 1983; Young and Sachs, 1979). These analyses provide sufficient spectrotemporal resolution for analyzing responses to stationary speech tokens like those used in many previous studies. To quantify power along dynamic formant trajectories in a nonstationary speech stimulus, the harmonicgram can be used as it offers superior spectrotemporal resolution compared to the spectrogram (Parida et al., 2021).

Briefly, the harmonicgram was constructed as follows. The fundamental frequency contour (F_0_[n], where *n* is the discrete index of time) was estimated from the sentence stimulus using Praat (Boersma, 2001). Response components (*HG*[*k, n*]) along the k^th^-harmonic of F_0_[n] were estimated using frequency demodulation of the difference PSTH *d*[*n*] followed by low-pass filtering (LPF). The low-pass filter was a 6^th^ order zero-phase IIR filter with 10-Hz cut-off frequency.

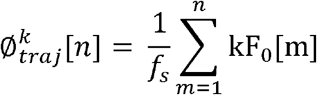

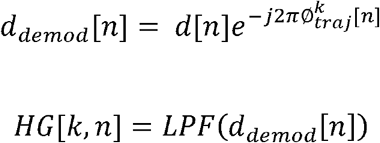

To evaluate the coding strength of a formant, F_x_, in the response d[n], the fractional power (*FracPower*) of three harmonics near F_0_-normalized F_x_ was calculated as:

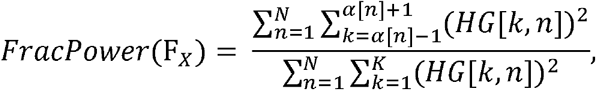

where 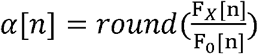, N is the length of d[n], *round* is the nearest-integer operator, and K is the total number of F_0_ harmonics considered. Because the lowest value for F_0_[n] was ~100 Hz and phase-locking for chinchillas is substantially degraded beyond 3500 Hz, K was set to 35 in these analyses.

For a given SNR condition, fractional power metrics were computed for responses to noisy-speech [*FracPower_SN_*(*F_x_*] and noise-alone [*FracPower_N_*(*F_x_*] for individual AN fibers. The difference in these two power metrics [i.e., *FracPower_SN_*(*F_x_*) – *FracPower_N_*(*F_x_*)] was used as the strength of *F_x_* coding for each fiber for that SNR condition. For the quiet condition, *FracPower_N_*(*F_x_*) was set to 0.

##### Coding fidelity metric for fricatives in noise based on correlation analysis

To evaluate the fidelity of the neural representation of the fricative /s/ in noise, correlations of the slowly varying envelope responses were quantified between speech-alone and noisy-speech conditions during the fricative segment. To confirm that these correlation values were not spurious correlations between speech and noise, the speech-alone and noisy-speech correlations were corrected by subtracting the correlation between response envelopes of speech-alone and noise-alone conditions for the same fricative window.

Response envelopes were obtained from single-polarity PSTHs using a low-pass filter (fourthorder, cut-off=32 Hz). Let the response envelope to speech-alone, noisy-speech, and noise-alone be denoted by *R_S_, R_SN_*, and *R_N_*, respectively. Then, a speech-in-noise coding fidelity metric was quantified by computing the corrected correlation between *R_s_* and *R_SN_* as:

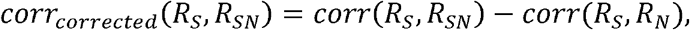

where *corr*(*X, Y*) = *E*(*XY*) and *E* is the expectation operator. Minimum value of *corr_corrected_*(*R_S_, R_SN_*) was set to 0.001 for display purposes in Figures 8D - 8F.

Plosive coding in noise was not considered because plosives were completely masked by noise, even at the highest SNR (i.e., 0 dB) used here. Instead, inferences about plosive coding in noise based on formant transitions (as quantifying in Figure 5) are considered in the *Discussion*.

##### Statistical analyses

Statistical analyses of group effects (i.e., hearing status) were performed in R (version 4.0.3) using linear mixed-effects models (lme4 package, Bates et al., 2014). Both p- and F-values were estimated using Type II Wald F tests (Kenward and Roger, 1997). Log-transformed CF and its interaction with hearing status were included in all statistical models for AN data. CF was log-transformed because of the logarithmic spacing of frequency in the cochlea. Appropriate CF ranges for statistical analyses were determined based on stimulus spectral bandwidth and are described in each figure caption. Driven rate and spontaneous rate were included as predictors in the temporal-precision analyses in Figure 2. Spontaneous-rate group (low and medium SR, <18 spikes/s; high SR, >18 spikes/s) was included as a binomial variable in Figure 8. Interactions were included between fixed effects, with main and interaction effects judged to be significant when p<0.05.

The effects of various physiological mechanistic factors on speech-coding fidelity in Figure 4 (tonotopic coding), Figure 5 (formant coding in noise), and Figure 8 (fricative coding in noise) were also evaluated by including the following fixed-effect predictors: log-transformed CF, FTC threshold (in dB SPL), TTR (in dB), and local Q10. Animal identifier was treated as a random effect. The d_equivalent_ or d_eq_ metric was used to indicate effect size of significant and close-to-significant individual factors, i.e., d_eq_ is reported for all effect where p < 0.1 (Rosenthal and Rubin, 2003).

## Notes

https://github.com/HeinzLabPurdue/DT_SpINcoding-paper

### Competing Interest Statement

The authors have declared no competing interest.

## REFERENCES

Auerbach, B.D., Rodrigues, P.V., and Salvi, R.J. (2014). Central Gain Control in Tinnitus and Hyperacusis. Front. Neurol. 5.

Bates, D., Mächler, M., Bolker, B., and Walker, S. (2014). Fitting linear mixed-effects models using lme4. ArXiv Prepr. ArXiv14065823.

Bharadwaj, H.M., Verhulst, S., Shaheen, L., Liberman, M.C., and Shinn-Cunningham, B.G. (2014). Cochlear neuropathy and the coding of supra-threshold sound. Front. Syst. Neurosci. 8.

Bilger, R.C., and Wang, M.D. (1976). Consonant confusions in patients with sensorineural hearing loss. J. Speech Hear. Res. 19, 718–748.

Boersma, P. (2001). Praat, a system for doing phonetics by computer. Glot Int 5, 341–345.

Chicharro, D., Kreuz, T., and Andrzejak, R.G. (2011). What can spike train distances tell us about the neural code? J. Neurosci. Methods 199, 146–165.

Chintanpalli, A., and Heinz, M.G. (2007). Effect of auditory-nerve response variability on estimates of tuning curves. J. Acoust. Soc. Am. 122, EL203–EL209.

Chung, K. (2004). Challenges and Recent Developments in Hearing Aids: Part I. Speech Understanding in Noise, Microphone Technologies and Noise Reduction Algorithms. Trends Amplif. 8, 83–124.

Clark, J.G. (1981). Uses and abuses of hearing loss classification. Asha 23, 493–500.

Cooke, M. (2006). A glimpsing model of speech perception in noise. J. Acoust. Soc. Am. 119, 1562–1573.

Costalupes, J.A., Young, E.D., and Gibson, D.J. (1984). Effects of continuous noise backgrounds on rate response of auditory nerve fibers in cat. J Neurophysiol 51, 1326–1344.

Dawes, P., Emsley, R., Cruickshanks, K.J., Moore, D.R., Fortnum, H., Edmondson-Jones, M., McCormack, A., and Munro, K.J. (2015). Hearing Loss and Cognition: The Role of Hearing Aids, Social Isolation and Depression. PLOS ONE 10, e0119616.

Delgutte, B., and Kiang, N.Y. (1984a). Speech coding in the auditory nerve: V. Vowels in background noise. J. Acoust. Soc. Am. 75, 908–918.

Delgutte, B., and Kiang, N.Y.S. (1984b). Speech coding in the auditory nerve: I. Vowel like sounds. J. Acoust. Soc. Am. 75, 866–878.

Delgutte, B., and Kiang, N.Y.S. (1984c). Speech coding in the auditory nerve: III. Voiceless fricative consonants. J. Acoust. Soc. Am. 75, 887–896.

Delgutte, B., Hammond, B.M., and Cariani, P.A. (1998). Neural coding of the temporal envelope of speech: relation to modulation transfer functions. Psychophys. Physiol. Adv. Hear. 595–603.

Dubno, J.R., and Dirks, D.D. (1989). Auditory filter characteristics and consonant recognition for hearing impaired listeners. J. Acoust. Soc. Am. 85, 1666–1675.

Dubno, J.R., Dirks, D.D., and Langhofer, L.R. (1982). Evaluation of hearing-impaired listeners using a nonsense-syllable test II. Syllable recognition and consonant confusion patterns. J. Speech Lang. Hear. Res. 25, 141–148.

Elhilali, M. (2019). Modulation Representations for Speech and Music. In Timbre: Acoustics, Perception, and Cognition, K. Siedenburg, C. Saitis, S. McAdams, A.N. Popper, and R.R. Fay, eds. (Cham: Springer International Publishing), pp. 335–359.

Elliott, T.M., and Theunissen, F.E. (2009). The Modulation Transfer Function for Speech Intelligibility. PLoS Comput Biol 5, e1000302.

Fernandez, K.A., Jeffers, P.W.C., Lall, K., Liberman, M.C., and Kujawa, S.G. (2015). Aging after Noise Exposure: Acceleration of Cochlear Synaptopathy in “Recovered” Ears. J. Neurosci. 35, 7509–7520.

Fernandez, K.A., Guo, D., Micucci, S., De Gruttola, V., Liberman, M.C., and Kujawa, S.G. (2020). Noise-induced Cochlear Synaptopathy with and Without Sensory Cell Loss. Neuroscience 427, 43–57.

Festen, J.M., and Plomp, R. (1983). Relations between auditory functions in impaired hearing. J. Acoust. Soc. Am. 73, 652–662.

Festen, J.M., and Plomp, R. (1990). Effects of fluctuating noise and interfering speech on the speech reception threshold for impaired and normal hearing. J. Acoust. Soc. Am. 88, 1725–1736.

French, N.R., and Steinberg, J.C. (1947). Factors Governing the Intelligibility of Speech Sounds. J. Acoust. Soc. Am. 19, 90–119.

Glasberg, B.R., and Moore, B.C. (1989). Psychoacoustic abilities of subjects with unilateral and bilateral cochlear hearing impairments and their relationship to the ability to understand speech. Scand. Audiol. Suppl. 32, 1–25.

Glasberg, B.R., and Moore, B.C.J. (1986). Auditory filter shapes in subjects with unilateral and bilateral cochlear impairments. J. Acoust. Soc. Am. 79, 1020–1033.

Goman, A.M., and Lin, F.R. (2016). Prevalence of Hearing Loss by Severity in the United States. Am. J. Public Health 106, 1820–1822.

Halliday, L.F., Rosen, S., Tuomainen, O., and Calcus, A. (2019). Impaired frequency selectivity and sensitivity to temporal fine structure, but not envelope cues, in children with mild-to-moderate sensorineural hearing loss. J. Acoust. Soc. Am. 146, 4299–4314.

Harrison, R.V., and Evans, E.F. (1979). Some aspects of temporal coding by single cochlear fibres from regions of cochlear hair cell degeneration in the guinea pig. Arch. Otorhinolaryngol. 224, 71–78.

Heinz, M.G., Issa, J.B., and Young, E.D. (2005). Auditory-Nerve Rate Responses are Inconsistent with Common Hypotheses for the Neural Correlates of Loudness Recruitment. J. Assoc. Res. Otolaryngol. 6, 91–105.

Henry, K.S., and Heinz, M.G. (2012). Diminished temporal coding with sensorineural hearing loss emerges in background noise. Nat. Neurosci. 15, 1362–1364.

Henry, K.S., Kale, S., Scheidt, R.E., and Heinz, M.G. (2011). Auditory brainstem responses predict auditory nerve fiber thresholds and frequency selectivity in hearing impaired chinchillas. Hear. Res. 280, 236–244.

Henry, K.S., Kale, S., and Heinz, M.G. (2016). Distorted Tonotopic Coding of Temporal Envelope and Fine Structure with Noise-Induced Hearing Loss. J. Neurosci. 36, 2227–2237.

Henry, K.S., Sayles, M., Hickox, A.E., and Heinz, M.G. (2019). Divergent auditory-nerve encoding deficits between two common etiologies of sensorineural hearing loss. J. Neurosci. 39, 6879--6887.

Hopkins, K., Moore, B.C.J., and Stone, M.A. (2008). Effects of moderate cochlear hearing loss on the ability to benefit from temporal fine structure information in speech. J Acoust Soc Am 123, 1140–1153.

Horst, J.W. (1987). Frequency discrimination of complex signals, frequency selectivity, and speech perception in hearing impaired subjects. J. Acoust. Soc. Am. 82, 874–885.

Kale, S., and Heinz, M.G. (2010). Envelope Coding in Auditory Nerve Fibers Following Noise-Induced Hearing Loss. J. Assoc. Res. Otolaryngol. 11, 657–673.

Kenward, M.G., and Roger, J.H. (1997). Small Sample Inference for Fixed Effects from Restricted Maximum Likelihood. Biometrics 53, 983–997.

Kiang, N.Y.S., and Moxon, E.C. (1974). Tails of tuning curves of auditory□nerve fibers. J. Acoust. Soc. Am. 55, 620–630.

King, A., Hopkins, K., and Plack, C.J. (2016). Differential Group Delay of the Frequency Following Response Measured Vertically and Horizontally. J. Assoc. Res. Otolaryngol. 17, 133–143.

Le Prell, C.G., and Clavier, O.H. (2017). Effects of noise on speech recognition: Challenges for communication by service members. Hear. Res. 349, 76–89.

Lesica, N.A. (2018). Why Do Hearing Aids Fail to Restore Normal Auditory Perception? Trends Neurosci.

Liberman, M.C. (1984). Single-neuron labeling and chronic cochlear pathology. I. Threshold shift and characteristic-frequency shift. Hear. Res. 16, 33–41.

Liberman, M.C., and Dodds, L.W. (1984a). Single-neuron labeling and chronic cochlear pathology. III. Stereocilia damage and alterations of threshold tuning curves. Hear. Res. 16, 55–74.

Liberman, M.C., and Dodds, L.W. (1984b). Single-neuron labeling and chronic cochlear pathology. II. Stereocilia damage and alterations of spontaneous discharge rates. Hear. Res. 16, 43–53.

Liberman, M.C., and Dodds, L.W. (1984c). Single-neuron labeling and chronic cochlear pathology. III. Stereocilia damage and alterations of threshold tuning curves. Hear Res 16, 55–74.

Lorenzi, C., Gilbert, G., Carn, H., Garnier, S., and Moore, B.C.J. (2006). Speech perception problems of the hearing impaired reflect inability to use temporal fine structure. Proc. Natl. Acad. Sci. 103, 18866–18869.

Lybarger, S.F. (1978). Selective amplification—a review and evaluation. Ear Hear. 3, 258.

McCormack, A., and Fortnum, H. (2013). Why do people fitted with hearing aids not wear them? Int. J. Audiol. 52, 360–368.

Mener, D.J., Betz, J., Genther, D.J., Chen, D., and Lin, F.R. (2013). Hearing Loss and Depression in Older Adults. J. Am. Geriatr. Soc. 61, 1627–1629.

Miller, R.L., Schilling, J.R., Franck, K.R., and Young, E.D. (1997). Effects of acoustic trauma on the representation of the vowel */ε/* in cat auditory nerve fibers. J. Acoust. Soc. Am. 101, 3602–3616.

Moore, B.C.J. (2007). Cochlear Hearing Loss: Physiological, Psychological, and Technical Issues, 2nd edition (Chichester: Wiley).

Moore, B.C.J. (2008). The role of temporal fine structure processing in pitch perception, masking, and speech perception for normal-hearing and hearing-impaired people. J Assoc Res Otolaryngol 9, 399–406.

Moore, B.C.J., Glasberg, B.R., Hess, R.F., and Birchall, J.P. (1985). Effects of flanking noise bands on the rate of growth of loudness of tones in normal and recruiting ears. J. Acoust. Soc. Am. 77, 1505–1513.

Ngan, E.M., and May, B.J. (2001). Relationship between the auditory brainstem response and auditory nerve thresholds in cats with hearing loss. Hear. Res. 156, 44–52.

Nielsen, J.B., and Dau, T. (2009). Development of a Danish speech intelligibility test. Int. J. Audiol. 48, 729–741.

Parida, S., and Heinz, M.G. (2021). Noninvasive Measures of Distorted Tonotopic Speech Coding Following Noise-Induced Hearing Loss. J. Assoc. Res. Otolaryngol. 22, 51–66.

Parida, S., Bharadwaj, H., and Heinz, M.G. (2021). Spectrally specific temporal analyses of spike-train responses to complex sounds: A unifying framework. PLOS Comput. Biol. 17, e1008155.

Phatak, S.A., and Grant, K.W. (2014). Phoneme recognition in vocoded maskers by normalhearing and aided hearing-impaired listeners. J. Acoust. Soc. Am. 136, 859–866.

Rosen, S. (1992). Temporal information in speech: acoustic, auditory, and linguistic aspects. Phil Trans R Soc Lond B 336, 367–373.

Rosenthal, R., and Rubin, D.B. (2003). r(equivalent:) A simple effect size indicator. Psychol. Methods 8, 492–496.

Ruggero, M.A., and Rich, N.C. (1991). Furosemide alters organ of Corti mechanics: Evidence for feedback of outer hair cells upon the basilar membrane. J Neurosci 11, 1057–1067.

Sachs, M.B., Voigt, H.F., and Young, E.D. (1983). Auditory nerve representation of vowels in background noise. J Neurophysiol 50, 27–45.

Sayles, M., and Heinz, M.G. (2017). Afferent Coding and Efferent Control in the Normal and Impaired Cochlea. In Understanding the Cochlea, G.A. Manley, A.W. Gummer, A.N. Popper, and R.R. Fay, eds. (Cham Switzerland: Springer), pp. 215–252.

Scheidt, R.E., Kale, S., and Heinz, M.G. (2010). Noise-induced hearing loss alters the temporal dynamics of auditory-nerve responses. Hear. Res. 269, 23–33.

Schilling, J.R., Miller, R.L., Sachs, M.B., and Young, E.D. (1998). Frequency-shaped amplification changes the neural representation of speech with noise-induced hearing loss. Hear. Res. 117, 57–70.

Silkes, S.M., and Geisler, C.D. (1991). Responses of ‘“lower□spontaneous □rate”‘ auditory□nerve fibers to speech syllables presented in noise. I: General characteristics. J. Acoust. Soc. Am. 90, 3122–3139.

Sinex, D.G., and Geisler, C.D. (1983). Responses of auditory□nerve fibers to consonant–vowel syllables. J. Acoust. Soc. Am. 73, 602–615.

Skoe, E., and Kraus, N. (2010). Auditory brainstem response to complex sounds: a tutorial. Ear Hear. 31, 302–324.

Strelcyk, O., and Dau, T. (2009). Relations between frequency selectivity, temporal fine-structure processing, and speech reception in impaired hearing. J. Acoust. Soc. Am. 125, 3328–3345.

Thomson, D.J. (1982). Spectrum estimation and harmonic analysis. Proc. IEEE 70, 1055–1096.

Trevino, M., Lobarinas, E., Maulden, A.C., and Heinz, M.G. (2019). The chinchilla animal model for hearing science and noise-induced hearing loss. J. Acoust. Soc. Am. 146, 3710–3732.

Turner, C.W., and Robb, M.P. (1987). Audibility and recognition of stop consonants in normal and hearing impaired subjects. J. Acoust. Soc. Am. 81, 1566–1573.

Van de Grift Turek, S., Dorman, M.F., Franks, J.R., and Summerfield, Q. (1980). Identification of synthetic /bdg/ by hearing impaired listeners under monotic and dichotic formant presentation. J. Acoust. Soc. Am. 67, 1031–1040.

Victor, J.D., and Purpura, K.P. (1996). Nature and precision of temporal coding in visual cortex: a metric-space analysis. J. Neurophysiol. 76, 1310–1326.

Woolf, N.K., Ryan, A.F., and Bone, R.C. (1981). Neural phase-locking properties in the absence of cochlear outer hair cells. Hear. Res. 4, 335–346.

Wu, P.Z., Liberman, L.D., Bennett, K., de Gruttola, V., O’Malley, J.T., and Liberman, M.C. (2019). Primary Neural Degeneration in the Human Cochlea: Evidence for Hidden Hearing Loss in the Aging Ear. Neuroscience 407, 8–20.

Young, E.D. (2008). Neural representation of spectral and temporal information in speech. Philos. Trans. R. Soc. B Biol. Sci. 363, 923–945.

Young, E.D. (2012). Neural Coding of Sound with Cochlear Damage. In Noise-Induced Hearing Loss: Scientific Advances, C. LePrell, and D. Henderson, eds. (New York: Springer), pp. 87–135.

Young, E.D., and Barta, P.E. (1986). Rate responses of auditory nerve fibers to tones in noise near masked threshold. J Acoust Soc Am 79, 426–442.

Young, E.D., and Sachs, M.B. (1979). Representation of steady□state vowels in the temporal aspects of the discharge patterns of populations of auditory□nerve fibers. J. Acoust. Soc. Am. 66, 1381–1403.

Zeng, F.G., and Turner, C.W. (1990). Recognition of voiceless fricatives by normal and hearing-impaired subjects. J. Speech Hear. Res. 33, 440–449.

Zeng, F.G., and Turner, C.W. (1991). Binaural loudness matches in unilaterally impaired listeners. Q J Exp Psychol A 43, 565–583.

Zhong, Z., Henry, K.S., and Heinz, M.G. (2014). Sensorineural hearing loss amplifies neural coding of envelope information in the central auditory system of chinchillas. Hear. Res. 309, 55–62.

